# A systematic evaluation of computational methods for cell segmentation

**DOI:** 10.1101/2024.01.28.577670

**Authors:** Yuxing Wang, Junhan Zhao, Hongye Xu, Cheng Han, Zhiqiang Tao, Dawei Zhou, Tong Geng, Dongfang Liu, Zhicheng Ji

## Abstract

Cell segmentation is a fundamental task in analyzing biomedical images. Many computational methods have been developed for cell segmentation and instance segmentation, but their performances are not well understood in various scenarios. We systematically evaluated the performance of 18 segmentation methods to perform cell nuclei and whole cell segmentation using light microscopy and fluorescence staining images. We found that general-purpose methods incorporating the attention mechanism exhibit the best overall performance. We identified various factors influencing segmentation performances, including image channels, choice of training data, and cell morphology, and evaluated the generalizability of methods across image modalities. We also provide guidelines for choosing the optimal segmentation methods in various real application scenarios. We developed Seggal, an online resource for downloading segmentation models already pre-trained with various tissue and cell types, substantially reducing the time and effort for training cell segmentation models.

## INTRODUCTION

Cell morphology, which describes the shape, size, and structure of cells, is crucial for understanding the types and states of cells and how cells respond to their microenvironments^1–7^. The study of cell morphology is enabled by biomedical imaging technologies such as phase-contrast microscopy^8,9^ and multiplexed fluorescence imaging^10^. However, these technologies do not directly provide the boundaries of cells, and the cell boundaries must be defined either manually or by computational methods. While manually identifying cell boundaries is infeasible for a large number of images, computational methods for cell segmentation become indispensable for delineating cell boundaries in most settings. Many downstream analysis tasks, such as modeling sub-cellular patterns^11,12^, screening genetic or chemical perturbations^13,14^, tracking the cell migration and development^15,16^, and deciphering cell physiological status^17,18^, depend on the computationally identified cell boundaries. Thus, an accurate computational method for cell segmentation is of paramount importance for appropriately handling biomedical images and extracting insights from the data.

The advancement of deep learning in the past decade has inspired numerous cell and instance segmentation methods that significantly outperform traditional approaches, such as the watershed method^19^. These methods can be categorized into two classes. The first class includes methods developed and tested specifically for biomedical images, such as Cellpose^20^, StarDist^21^, RetinaMask^22^, Mesmer^23^, and FeatureNet^24^. The second class comprises methods like Centermask2^25^, Mask RCNN^26^, ResNeSt^27^, MS RCNN^28^, Mask2former^29^, Cascade Mask RCNN seesaw^30^, Swin Transformer^31^, SOLOv2^32^, RF-Next^33^, HRNet^34^, Res2Net^35^, and Segment Anything^36^, developed for more general purposes of instance segmentation. In principle, these general-purpose methods are applicable to biomedical images for cell segmentation, as it is a form of instance segmentation. However, they were primarily tested on generic datasets such as ImageNet^37^, and their effectiveness in cell segmentation remains largely unexplored.

The performances of cell segmentation methods were compared in previous studies, including the Cell Tracking Challenge^38^, the 2018 Data Science Bowl challenge^39^, and a recent grand challenge organized by NeurIPS^40^. However, these studies have several limitations. First, the volumn of data included in these studies is limited. For instance, the 2018 Data Science Bowl provides 37,333 annotated nuclei, and the NeurIPS challenge provides 168,491 annotated cells. These numbers are substantially smaller than those of recently published datasets like LIVECell^8,41^ and TissueNet^10,42^, each containing more than a million annotated cells or nuclei. Since these large datasets are publicly available for training cell segmentation methods, the results of previous challenges no longer reflect the state-of-the-art performance of cell segmentation methods. Second, the performance of cell segmentation methods can vary dramatically across tissue types and imaging technologies. Previous benchmark studies focus only on the overall performance, averaged across various scenarios, and lack detailed evaluations for each tissue type and imaging technology. Since most segmentation tasks involve one tissue type using one imaging technology, the value of previous benchmark studies is limited in real-world practice for selecting the most appropriate segmentation method. Third, studies like the NeurIPS and Data Science Bowl challenges focus on evaluating methods submitted by challenge participants. These methods often lack open-source software packages with well-documented vignettes, making them difficult to apply to new datasets, especially for users without extensive training in computer programming and machine learning. In summary, a more systematic evaluation of cell segmentation methods is still needed to guide the practice of cell segmentation in real-world applications.

To address these limitations, we conducted a systematic evaluation of computational methods for cell segmentation, aiming to provide guidelines for applying and developing these methods in practical applications. The importance of this study is threefold. For researchers who are designing experiments, this study can help to decide a cost-effective strategy for generating imaging data. For researchers who have collected imaging data, this study can help identify the best-performing method and associated training data, depending on the available computational resources. For machine learning researchers, this study sheds light on potential opportunities that could lead to the development of more powerful cell segmentation methods in the future. In addition, we have also provided the pre-trained models for each cell segmentation method, tissue or cell type, and imaging technology as a publicly available and freely accessible resource, significantly reducing the effort and computational resources required for training cell segmentation methods.

## RESULTS

### Overview

In this study, we evaluated the performance of 18 instance segmentation methods for both light microscopy and fluorescence staining images (Supplementary Information). These 18 methods can be divided into two categories. The first category comprises methods designed for biomedical images, including Cellpose^20^, Mesmer^23^, StarDist^21^, RetinaMask^22^, and FeatureNet^24^. The second category comprises methods designed for generic instance segmentation tasks, including Centermask2^25^, Mask RCNN^26^, ResNeSt^27^, MS RCNN^28^, Mask2former^29^, Cascade Mask RCNN Seesaw^30^, Swin Transformer (Swin-S and Swin-T)^31^, SOLOv2^32^, RF-Next^33^, HRNet^34^, Res2Net^35^, and Segment Anything^36^. We include both Swin-S and Swin-T, two versions of the Swin Transformer with different model sizes, to explore how model complexity impacts segmentation performance. Segment Anything is based on a foundation model that does not require additional training or fine-tuning.

The entire evaluation dataset comprises more than 12,400 images and approximately 3 million cells, representing ten different tissue types and eight different cell types. We collected 7,022 images generated by multiplexed fluorescence imaging platforms from TissueNet^10,42^. Each image includes a nuclear channel (such as DAPI) and a membrane or cytoplasm channel (such as E-cadherin or Pan-Keratin). These images feature 1.3 million whole cells and 1.2 million cell nuclei from ten tissue types (Figure 1a). Additionally, we collected 5,387 images generated by phase-contrast microscopy from LIVECell^8,41^. These images are single-channel and contain 1,686,352 cells from eight cell types (Figure 1b). Both TissueNet and LIVECell provide annotated cell or nuclei masks for each image, which were used as training labels and as gold standards for evaluations.

**Figure 1.**
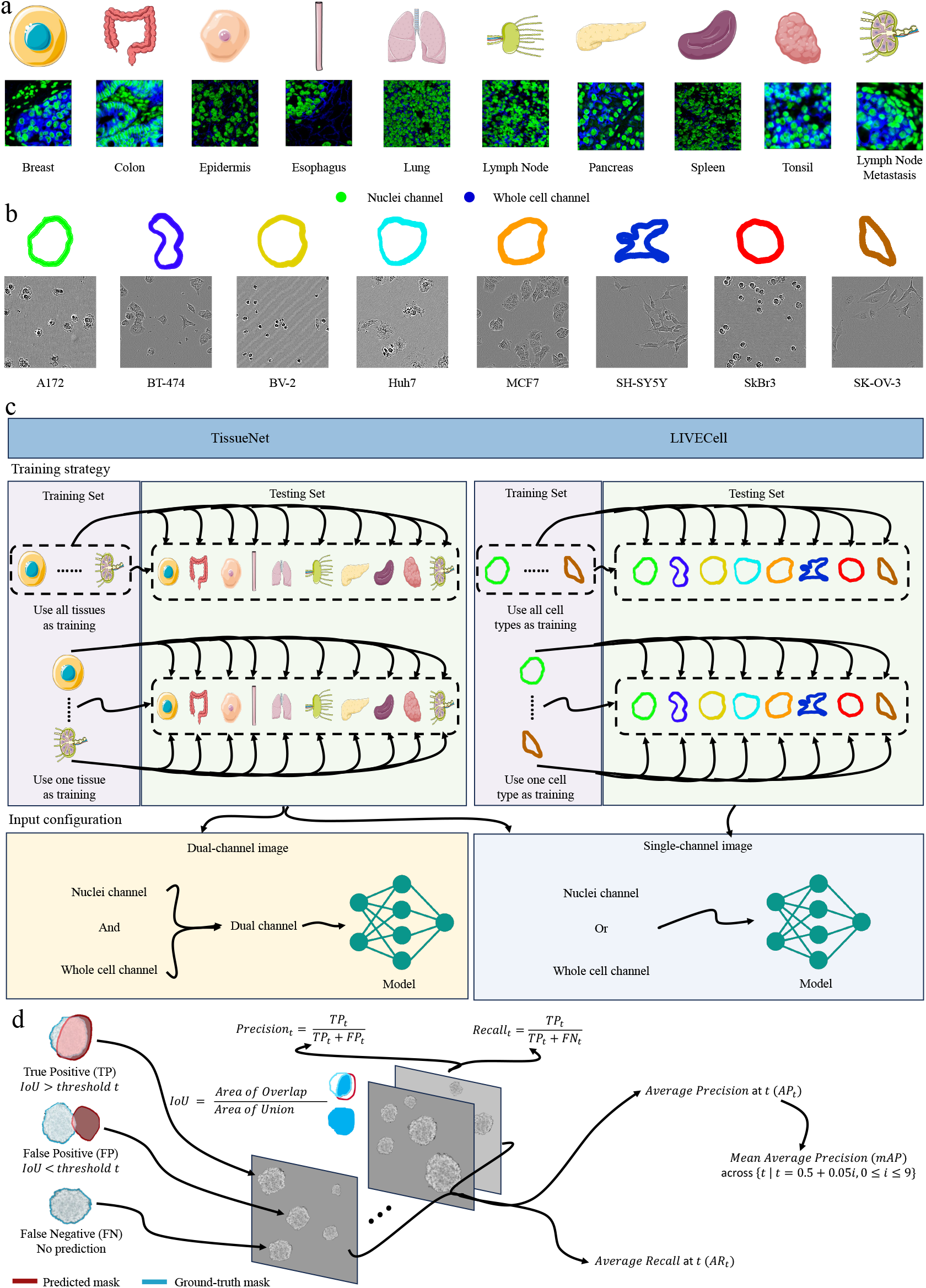
Overview of the whole study. **a**, Example fluorescence staining images from TissueNet for ten distinct tissue types. The green and blue channels represent cell nuclei and whole cells, respectively. **b**, Example microscopy images from LIVECell for eight distinct cell types. **c**, Segmentation models were trained on all tissue and cell types, as well as on individual tissues and cell types. For TissueNet, both dual-channel and single-channel images were used for segmenting cell nuclei and whole cells. For LIVECell, whole cell images were used for whole cell segmentation. **d**, The accuracy of cell segmentation methods was evaluated based on average precision, average recall, and mean average precision.

Five types of segmentation tasks were evaluated (Figure 1c). In TissueNet, segmentation methods were trained either with nuclear channel images or dual-channel images for cell nuclei segmentation. Similarly, methods were trained either with whole cell channel images or dual-channel images for whole cell segmentation. For LIVECell, both model training and whole cell segmentation were performed using whole cell channel images. We assessed the performance of segmentation methods trained by aggregating images from all tissue types or cell types, as well as those trained with images from a single tissue type or cell type. The performance of each method was evaluated by comparing the predicted cell boundaries to the gold standard cell masks (Figure 1d, Methods). In accordance with the COCO evaluation metrics^23,43,44^, we employed multiple quantitative metrics, including average precision (AP), mean average precision (mAP), and average recall (AR).

In addition to segmentation accuracy, we also assessed the scalability and usability of these methods. Scalability is measured by the running time of a method (Supplementary Information). Usability, gauged by code maintenance, ease of use, and hardware support (Supplementary Information), indicates the ease with which the method can be implemented and executed in a new computing environment. These metrics provide valuable insights for practical applications.

### General-purpose methods with attention mechanism have the best performance

We systematically evaluated the accuracy, scalability, and usability of 18 cell segmentation methods and ranked them based on overall performance (Figure 2a, Supplementary Figure 1, Methods). Figure 2b shows examples of cell nuclei segmentation with dual-channel images for each method. While the top-performing methods produce results highly consistent with the gold standard, underperforming methods exhibit various issues (Figure 2c). These include oversegmentation, where one cell is incorrectly segmented into multiple cells; undersegmentation, where multiple cells are incorrectly grouped into one cell; missing cells, where models fail to identify cells; and incorrectly identifying non-cellular structures as cells. MS RCNN fails to generate reasonable cell boundaries due to the vanishing gradient problem encountered during the training process.

**Figure 2.**
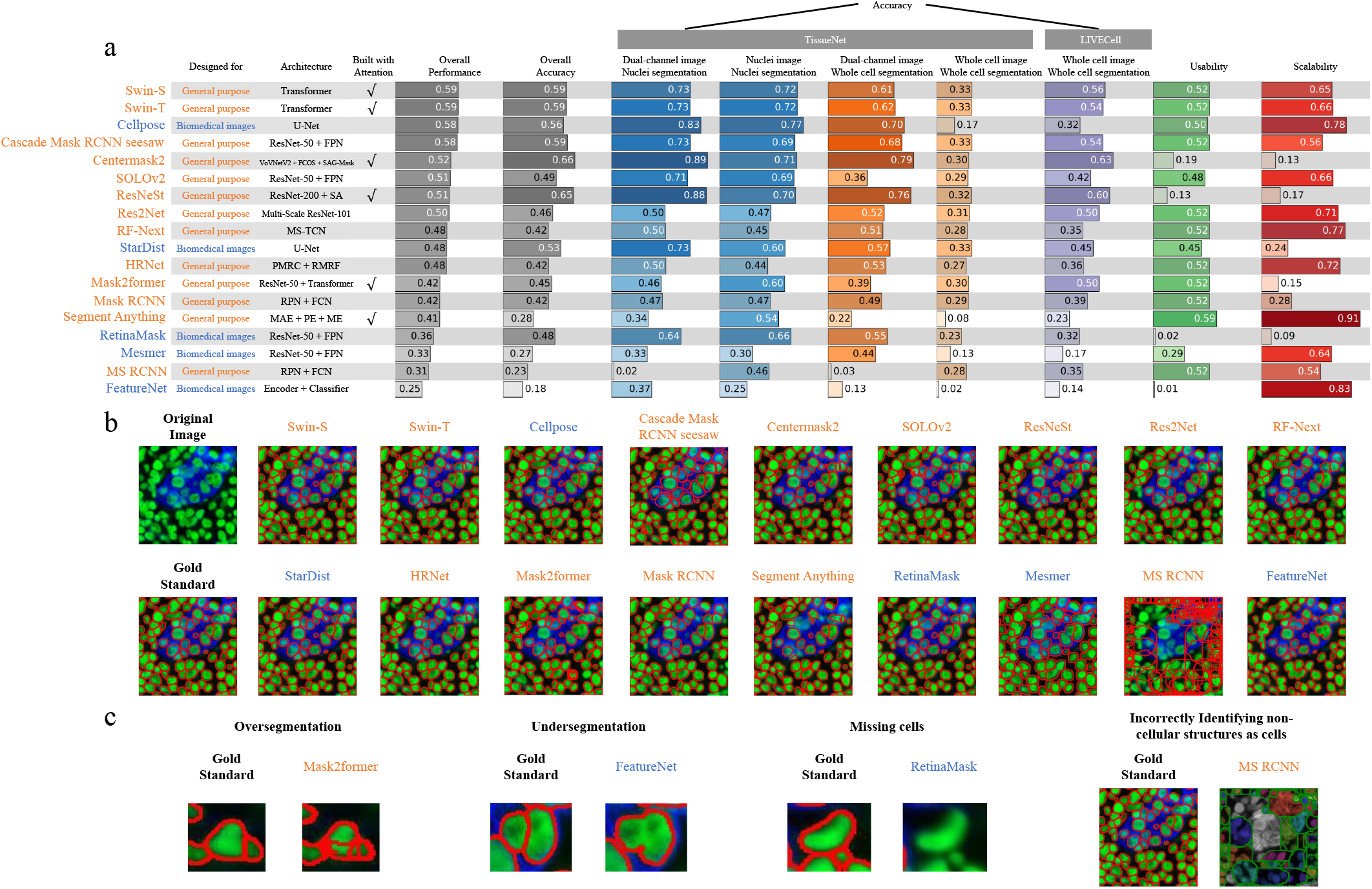
Overall performance of segmentation methods. **a**, Segmentation methods are ordered based on their overall performance. The overall performance is the average of five accuracy metrics along with usability and scalability. The overall accuracy is the average of five accuracy metrics. **b**, Examples of dual-channel images from TissueNet lymph node metastasis, including the original image, gold standard nuclei masks, and cell nuclei segmentation results of different methods. **c**, Examples of segmentation issues: oversegmentation, undersegmentation, missing cells, and incorrectly identifying non-cellular structures as cells.

Methods designed for general purposes outperform those developed specifically for biomedical images. Swin-S and Swin-T, both variants of the Swin Transformer^31^ and intended for general purposes, demonstrate the best overall performance. In each of the five accuracy metrics, both Swin-S and Swin-T consistently rank among the top 5 or 6 methods, evidencing their reliable performance across various scenarios. The two models exhibit almost identical performances, indicating that model complexity has minimal impact on the results for the Swin Transformer. Centermask2 and ResNeSt, also designed for general purposes, have the highest overall accuracy. In addition, they achieve the highest accuracy in segmenting both nuclei and whole cells using dual-channel images in the TissueNet dataset, and in segmenting whole cells in the LIVECell dataset. However, their overall performance is negatively impacted by their lower scalability and usability. Among methods specifically designed for biomedical imaging, Cellpose has the best overall performance. It shows the highest accuracy in segmenting nuclei with single-channel images in the TissueNet dataset and performs strongly whenever nuclei images are available. However, its performance significantly declines when nuclei images are absent. For whole cell segmentation with whole cell images, Centermask2, the best-performing general-purpose method, achieves improvements in accuracy of 96.9% and 76.5% over Cellpose in the LIVECell and TissueNet datasets, respectively. For whole cell segmentation with dual-channel images, Centermask2 achieves a 12.9% improvement in accuracy over Cellpose. Other methods tailored for biomedical images do not exhibit strong performance.

The general-purpose methods exhibiting the highest overall accuracy, such as Centermask2, ResNeSt, Swin-S, and Swin-T, all incorporate the attention mechanism^45^. In comparison, methods designed for biomedical imaging use architectures such as U-Net^46^ and ResNet^47^, which do not rely on the attention mechanism. Therefore, the use of the attention mechanism may explain why some general-purpose methods outperform those specifically designed for biomedical images. A potential reason for this improvement is that the attention mechanism allows the model to dynamically focus on the most relevant parts of an image, making it more adaptable to variations in biological images and leading to more accurate and efficient segmentation. While Mask2former and Segment Anything also use the attention mechanism, they underperform in this study. Notably, Segment Anything, a pre-trained foundation model not fine-tuned on our training images, may underperform due to this reason. Other methods employing different architectures, like Cascade Mask RCNN seesaw with ResNet^47^ and FPN^48^, also demonstrate competitive performance.

### Dual-channel images greatly enhance whole cell segmentation over single-channel images

Figure 2a compares the performance of segmentation methods trained with dual-channel versus single-channel images. Dual-channel images significantly enhance whole cell segmentation for most methods. For instance, Cellpose and Centermask2’s accuracy rose from 0.17 to 0.7 (312% improvement) and 0.3 to 0.79 (163% improvement), respectively. In contrast, the performance boost from dual-channel images is minimal for cell nuclei segmentation. Only a few methods like Centermask2, ResNeSt, and StarDist see substantial gains with over 0.1 accuracy increase. Methods such as Swin-S, SOLOv2, and Res2Net show similar performance with both image types. However, for some, including Mask2former and Segment Anything, performance notably declines with dual-channel images. These findings indicate varying capabilities among methods in integrating and utilizing multi-channel images. Additionally, while cell nuclei images alone suffice for nuclei segmentation, whole cell segmentation benefits from additional data in dual-channel images.

The performance gain from using dual-channel over single-channel images varies across different tissues (Supplementary Figure 2). For instance, in lung tissue, Centermask2’s accuracy for cell nuclei segmentation jumps from 0.308 to 0.655 (113% improvement), while in esophagus, the increase is more modest, from 0.691 to 0.759 (9.8% improvement). These findings are crucial for experimental design. In tissues like the esophagus, using single-channel images for cell nuclei segmentation yields results comparable to dual-channel images, potentially reducing experiment costs. However, for whole cell segmentation and for cell nuclei segmentation in tissues like the lung, employing dual-channel images is advised for significantly better segmentation outcomes.

### Segmentation performance is influenced by training sample size and cell morphology

We explored the influence of various factors on cell segmentation performance across five tasks, using training images from all tissue or cell types. Figure 3a shows the mAP of different methods when tested on various tissue or cell types. Generally, there is a consistent trend in method performance across these tissues or cell types. For instance, lymph node metastasis, pancreas, and esophagus consistently yield reliable nuclei segmentation results with most methods. In contrast, lung and spleen are often associated with poor segmentation, primarily due to the extremely limited number of images in the training data (Supplementary Figure 3). These observations suggest that there are some common factors influencing segmentation performance across different methods.

**Figure 3.**
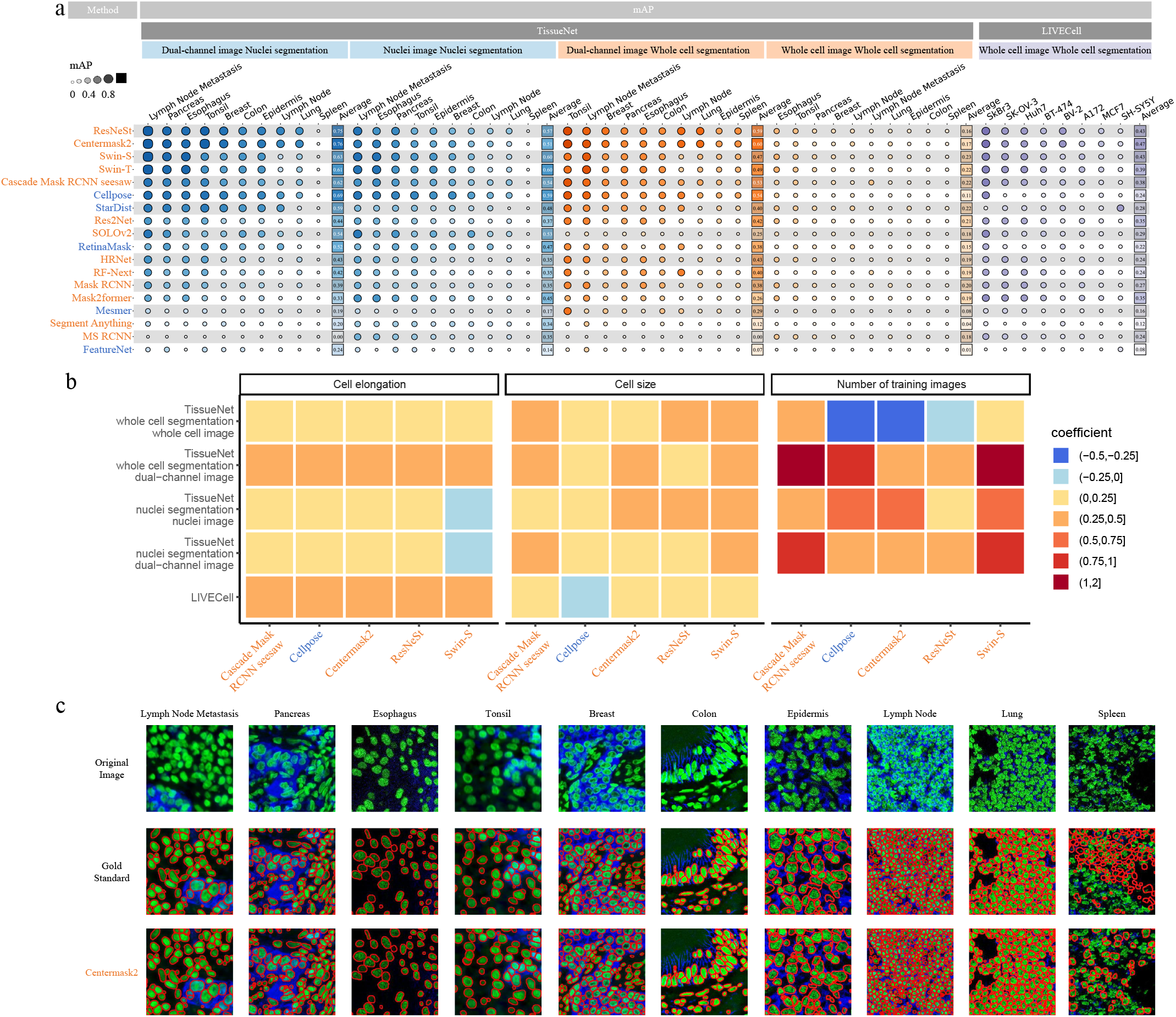
Segmentation performance with models trained on all tissues and cell types. **a**, mAP for each method (row) and for each tissue or cell type (column). Both rows and columns are ordered based on the average mAP. **b**, Coefficients of cell elongation, cell size, and number of training images in the multiple linear regression for mAP. The five methods with the highest overall accuracy are shown. **c**, Examples of dual-channel images from various TissueNet tissue types, including the original images, gold standard nuclei masks, and Centermask2 cell nuclei segmentation results.

To identify such common factors, we excluded lung and spleen from the TissueNet dataset and focused on tissue and cell types with sufficiently large numbers of training images. We employed a multiple linear regression model, considering the method’s mAP as the dependent variable, and the number of training images, cell elongation, and cell size as independent variables (Supplementary Information). Cell convexity was excluded due to its minimal variation across tissues (Supplementary Figure 4). For most methods and segmentation tasks, there is a positive correlation between segmentation performance and cell elongation, cell size, and the number of tissue-specific training images (Figure 3b). The correlation with the number of training images is expected, as images from the same tissue typically bear more resemblance to each other than to those from different tissues. The positive association with cell elongation and size indicates that segmentation algorithms are most effective for larger or rounder cells, such as those in lymph node metastasis. Conversely, smaller or irregularly shaped cells, like those in lymph nodes and the colon (Figure 3c, Supplementary Figure 5), pose challenges. This could be because smaller cells lack distinctive features for differentiation from the background, and irregularly shaped cells are less represented in the training data. Enhancing segmentation for such cells might be achieved by increasing their representation in the training set or integrating a model specifically trained on these cell types into the full model using an ensemble approach.

### Choice of training data impacts cell segmentation performance

We subsequently assessed how the performance of Swin-S, the top-performing method, is affected when trained with different tissue or cell types. Figures 4a-e and Supplementary Figures 6-9 illustrate Swin-S’s performance in the TissueNet dataset, trained individually with different tissue types or with all tissues combined. Notably, the segmentation performance is poorest when Swin-S is trained solely with lung or spleen images, likely due to their very limited number of training images. For tissues with adequate image quantities, segmentation performance varies with the training tissue. The best results are typically achieved when the same tissue is used for both training and segmentation. When training and segmentation tissues differ, some tissues demonstrate better transferability. For instance, Swin-S trained on breast tissue images consistently shows reliable nuclei segmentation across various tissues, whereas training with tonsil images results in subpar performance in tissues like the esophagus (Figure 4a). Furthermore, Swin-S trained on all tissues generally outperforms its counterpart trained on a single tissue, although the performance improvement is slight when the training and segmentation are on the same tissue. Figure 4f and Supplementary Figure 10 illustrate Swin-S’s performance when trained with individual versus all cell types in LIVECell, leading to conclusions similar to those from the TissueNet analysis. For instance, cell types like MCF-7 and BT-474 demonstrate relatively high transferability across different cell types.

**Figure 4.**
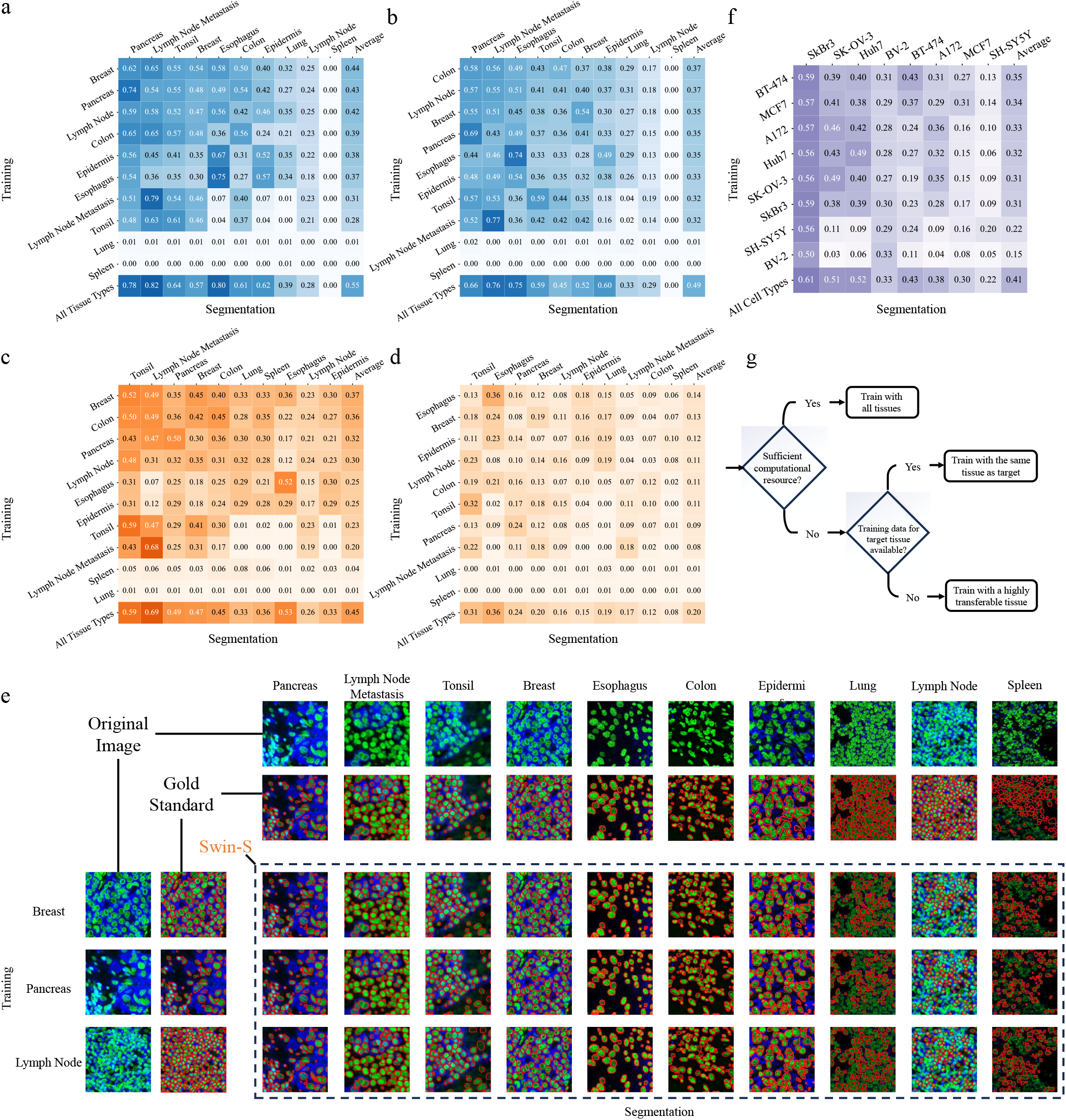
Segmentation performance with models trained on individual tissue and cell types. **a-d**, mAP of Swin-S with training (rows) and segmentation (columns) performed on individual TissueNet tissue types. mAP of Swin-S trained with all tissue types is also included as a reference. **a**: nuclei segmentation with dual-channel images. **b**: nuclei segmentation with nuclei images. **c**: whole cell segmentation with dual-channel images. **d**: whole cell segmentation with whole cell images. **e**, Examples of dual-channel images from various TissueNet tissue types for training (rows) and cell nuclei segmentation (columns), including the original images, gold standard nuclei masks, and Swin-S cell nuclei segmentation results. **f**, mAP of Swin-S with training (rows) and segmentation (columns) performed on individual LIVECell cell types. mAP of Swin-S trained with all cell types is also included as a reference. **g**, Strategy of choosing training data in real practices.

These results indicate that including images from diverse tissues and cell types in the training dataset is advantageous when sufficient computational resources are available. In scenarios with limited computational resources, it’s essential to determine the availability of training images from the same tissue or cell type as the target tissue for segmentation. Using such specific images for training often yields performance nearly equivalent to that achieved with a large, varied dataset. If not available, training should utilize images from tissues and cell types with high transferability, like breast tissue and MCF-7 cells. Figure 4g summarizes this practical strategy for selecting training data.

### Generalization of segmentation models

In real applications, the images employed for model training and segmentation often come from different modalities. For instance, *in situ* spatial transcriptomics data from platforms like Xenium typically provide only cell nuclei images, yet whole cell segmentation is necessary to fully capture a cell’s gene expression profile. Similarly, there are scenarios where only cell nuclei images are available for training, but segmentation must be conducted on whole cell images. These situations necessitate a segmentation method’s ability to generalize across multiple image modalities.

Figure 5a and Supplementary Figure 11a illustrate the performance of segmentation methods when training and segmentation use the same or different image modalities. The performance of most methods drops significantly when trained on whole cell images and applied to cell nuclei images, compared to training and segmenting on cell nuclei images. However, Cascade Mask RCNN seesaw and RF-Next maintain nearly identical performance in both scenarios, demonstrating strong generalizability across image modalities. A potential reason is the unique loss function and model architecture used by the two models. Cascade Mask RCNN seesaw introduces a novel seesaw loss function to dynamically re-balance the gradient issue between classes with abundant instances (head classes) and classes with few instances (tail classes). A few cells with almost identical cell and nuclei boundaries can be considered as tail classes in whole cell images. Cascade Mask RCNN seesaw can efficiently extract the features of these tail classes from whole cell images to be used in nuclei segmentation. RF-Next incorporates a global-to-local search strategy that efficiently and adaptively adjust and fine-tune receptive field combinations according to the specific characteristics of the data, leading to enhanced generalizability across different types of images. By contrast, all methods completely fail when trained on cell nuclei images and applied to whole cell images. These findings indicate that most segmentation methods struggle with generalizability across different image modalities.

**Figure 5.**
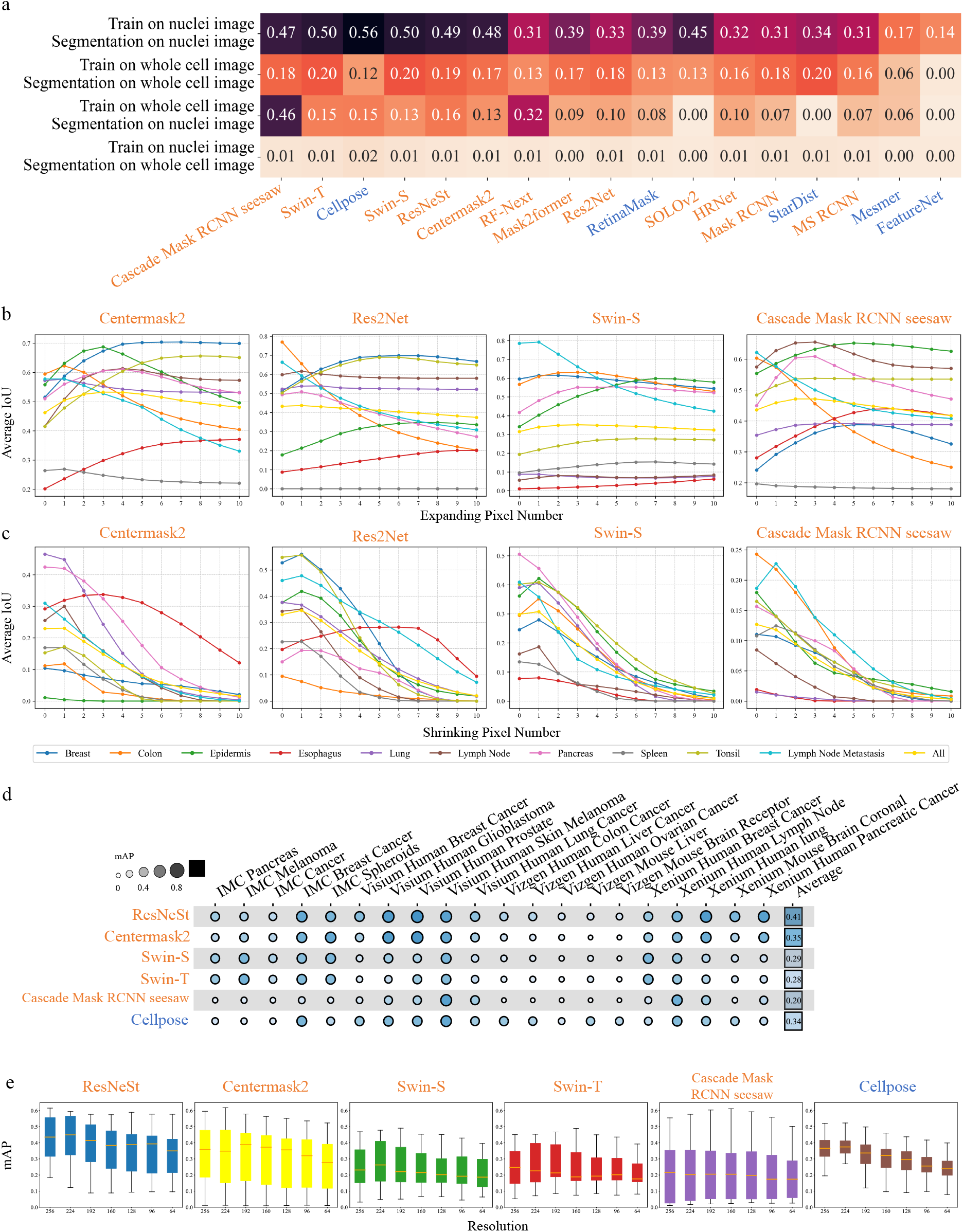
Generalizability across image modalities. **a**, Averaged mAP across TissueNet tissue types when training and segmentation were performed on the same or different image modalities. **b**, IoU (y-axis) between gold standard whole cell masks and expanded cell nuclei masks, averaged across cells within each TissueNet tissue type, when cell nuclei masks were expanded by different pixel numbers (x-axis). Zero expansion represents original cell nuclei masks without expansion. Four methods with the highest overall accuracy are shown. **c**, IoU (y-axis) between gold standard cell nuclei masks and shrunk whole cell masks, averaged across cells within each TissueNet tissue type, when whole cell masks were shrunk by different pixel numbers (x-axis). Zero shrinkage represents original whole cell masks without shrinkage. Four methods with the highest overall accuracy are shown. d. mAP scores of six top-performing methods across four technology platforms. e. mAP score (y-axis) and image resolution (x-axis) for each of the six top-performing methods.

We then explored a different strategy of segmenting images using the same modality as the training data, followed by morphologically dilating cell nuclei segmentation to approximate whole cell segmentation, or eroding whole cell segmentation for cell nuclei segmentation (Supplementary Information). This approach is currently used by 10x Xenium to provide an approximate whole cell segmentation, where the cell nuclei mask is expanded by 15 *μ*m or until reaching another cell boundary. Figure 5b and Supplementary Figure 11b demonstrate that expanding cell nuclei segmentation by varying pixel counts generally enhances whole cell segmentation performance. For example, in epidermis tissue, Swin-S shows a dramatic increase from 0.341 to 0.598 (75% improvement) with expansion. However, the optimal expansion degree varies across tissues. In ResNeSt, tonsil segmentation peaks at a 6-pixel expansion, whereas colon segmentation deteriorates with any nuclei boundary expansion. Therefore, a fixed-pixel expansion, as implemented by 10x Xenium, may not be optimal for all tissues. Figure 5c and Supplementary Figure 11c present the results of eroding whole cell segmentation for cell nuclei segmentation. The performance gains are more modest, with most cases showing optimal results with a one-pixel erosion.

Next, we evaluated the generalizability of methods across technology platforms. We applied six top-performing segmentation methods trained on TissueNet cell nuclei images to images generated by different technology platforms for nuclei segmentation. These platforms include imaging mass cytometry (IMC) and spatial transcriptomics (10x Visium, 10x Xenium, and Vizgen). Images from five tissues were included for each platform. For evaluation purposes, we manually annotated the nuclei boundaries of 100 cells for each tissue and platform and calculated mAP between the manually annotated and computationally generated nuclei boundaries (Figure 5d, Supplementary Figure 12). ResNeSt and Centermask2 are again top-performing methods. ResNeSt shows strong performances on 10x Visium and 10x Xenium, reaching an average mAP of 0.52 and 0.49, comparable to the performance in cross-validation studies (Figure 3a). In comparison, its performance is relatively weak on IMC and Vizgen datasets. These results suggest strong generalizability of top-performing methods across certain technology platforms.

Finally, we assessed the impact of image resolution on segmentation performance. Focusing on the three spatial transcriptomics platforms, we downsampled the original 256×256 images by factors of 0.125 to obtain images with six different resolutions: 224×224, 192×192, 160×160, 128×128, 96×96, and 64×64. We applied six top-performing methods to the original and downsampled images and evaluated their performances using mAP scores (Figure 5e). While there is an overall decreasing trend in performance with decreasing image resolutions, mAP scores of most methods remain high even for images with 64×64 resolution. These results suggest that the segmentation methods can be readily applied to images with lower resolutions.

### SegGal: a gallery of pre-trained cell segmentation models

As shown in this study (Figure 4) and in our previous work^7^, cell segmentation performance is significantly influenced by the training data. Pre-trained models, developed using a vast array of images from various tissues and cell types, can greatly enhance cell segmentation. However, training these models demands considerable computational resources and the effort to collect a large number of images. Despite their potential benefits, such comprehensive pre-trained models are often not readily available. While generic segmentation methods typically lack pre-trained models specialized for cell images, biomedical image segmentation methods like Cellpose only offer pre-trained models trained on a small numer of images.

To address this challenge, we have developed SegGal, a gallery of pre-trained cell segmentation models, leveraging the numerous models trained in this study (Figure 6a). SegGal includes 138 pre-trained models across 18 segmentation methods, catering to five different segmentation tasks. The training process utilized 450 GPUs and required 18,000GB of memory, spanning approximately 60 days. These pre-trained models are readily available for download from the SegGal website, allowing users to apply them directly to segmentation tasks. SegGal also provides detailed usage guides for installing and running various segmentation methods. This not only significantly reduces the computational load for model training but also streamlines the cell segmentation process.

**Figure 6.**
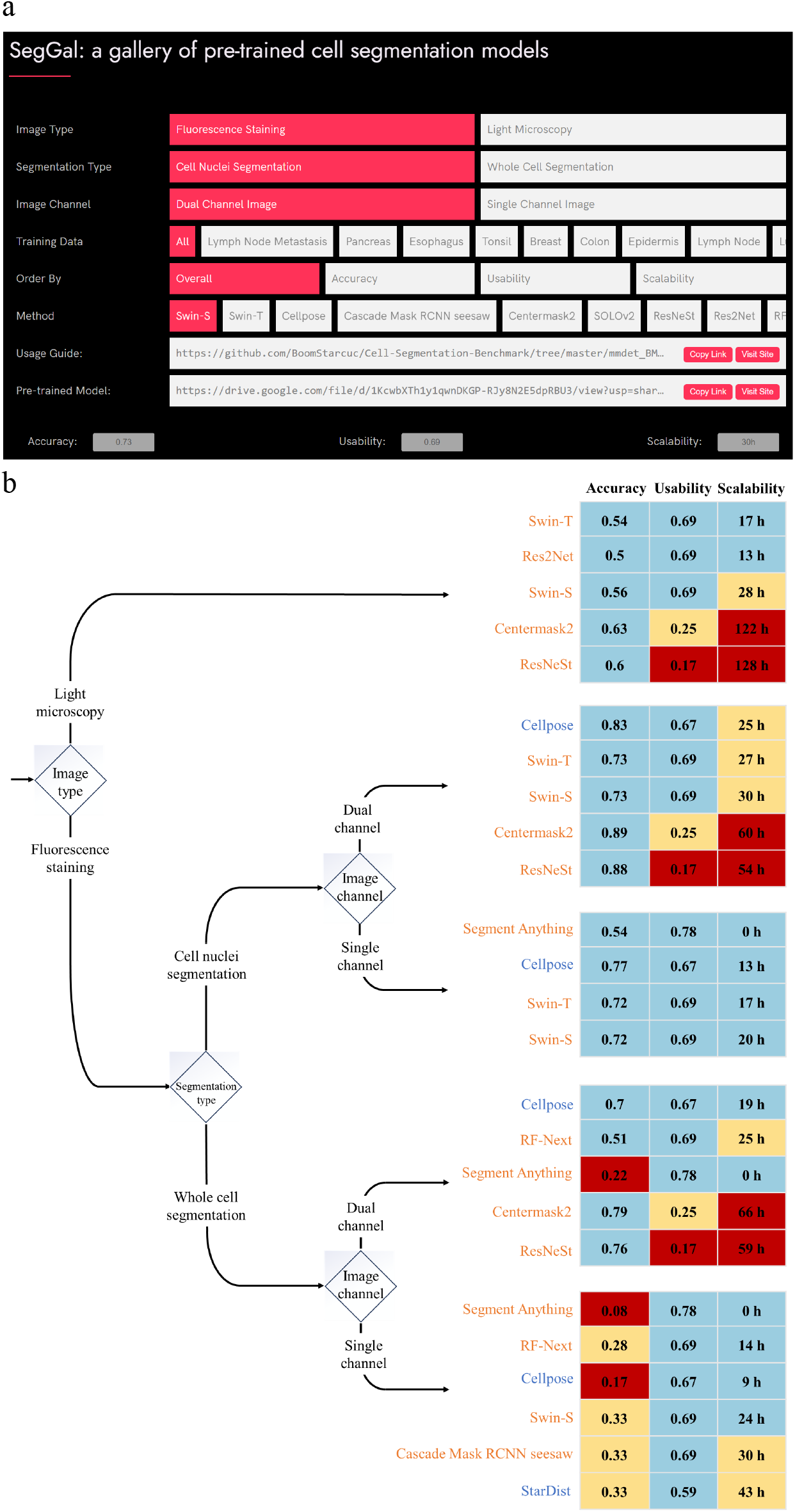
Applying cell segmentation methods in real practice. **a**, A screenshot of SegGal, an online resource that provides pre-trained models and usage guides for different segmentation methods under various scenarios. **b**, Guidelines for choosing the segmentation methods. Five different scenarios are listed based on image types, type of segmentation tasks, and availability of single-channel or dual-channel images. In each scenario, three methods with the best overall performance and three methods with the best overall accuracy were selected, and the union set of the six methods is shown. The methods are further ordered based on the overall performance. For accuracy and usability, values larger than or equal to 0.5 are shown as blue, values between 0.25 and 0.5 are shown as yellow, and values less than 0.25 are shown as red. For scalability, values less than or equal to 24 hours (h) are shown as blue, values between 24 h and 48 h are shown as yellow, and values larger than 48 h are shown as red.

## DISCUSSION

In this study, we systematically evaluated 18 segmentation methods for their effectiveness in segmenting cell nuclei and whole cells using multiplexed fluorescence and phase-contrast microscopy images from the TissueNet and LIVECell databases. These methods, encompassing both general-purpose and biomedical image-focused approaches, were assessed based on accuracy, usability, and scalability. We investigated various factors influencing segmentation performance, such as the use of single or dual-channel images, training sample size, choice of training data, and cell morphology. Additionally, we evaluated the methods’ ability to generalize across different image modalities. Lastly, we developed Seggal, an online resource providing access to a vast collection of pre-trained cell segmentation models.

Our findings contribute to the study of cell segmentation in three key areas. For researchers planning to generate imaging data, our study indicates that cell nuclei images are crucial for both cell nuclei and whole cell segmentation, with whole cell images proving beneficial in certain scenarios. For those developing cell segmentation methods, our findings highlight the attention mechanism, utilized by several top-performing methods, as a promising architecture worthy of further exploration. Additionally, our study offers a systematic guideline for researchers seeking to apply these methods in cell segmentation. Figure 6b provides a comprehensive list of cell segmentation methods suitable for various scenarios. Through the Seggal online resource, users gain direct access to a vast array of pre-trained models, significantly easing the burden of model training across different scenarios.

As indicated by this study, segmentation performance improves when training data includes a large variety of images from different tissues and cell types. Although Segment Anything, the sole foundation model evaluated in our study, did not perform well, we hypothesize that such a model could have significant potential if trained with massive, diverse datasets of cell images. TissueNet’s adoption of a human-in-the-loop approach demonstrates a viable method for generating extensive training datasets at a relatively low cost. Such datasets could pave the way for developing a foundational model broadly applicable to cell segmentation and potentially other biomedical image segmentation tasks.

## DATA and CODE AVAILABILITY

The original TissueNet dataset^42^ is available on the website: https://datasets.deepcell.org/data. The original LIVECell dataset^41^ is available on the website: https://sartorius-research.github.io/LIVECell/. The TissueNet dataset preprocessed in this study is available on Google Drive: https://drive.google.com/drive/folders/1dUtqhvkF-M7nSwtxUpY0QmgHIgA4pinc?usp=drive_link. The LIVECell dataset preprocessed in this study is available on Google Drive: https://drive.google.com/drive/folders/1mJayXI2W9DLL17fsD3j2AcFySebnsoza?usp=drive_link. Imaging Mass Cytometry (IMC) images were obtained from the `imcdatasets` R/Bioconductor package. 10x Visium and Xenium images were obtained from 10x website: https://www.10xgenomics.com/datasets.

Vizgen images were obtained from Vizgen website: https://info.vizgen.com. The SegGal online resource is available on the website: https://boomstarcuc.github.io/SegGal/. Software code used to perform the analysis of this study is available on GitHub: https://github.com/BoomStarcuc/Cell-Segmentation-Benchmark with License MIT.

## ACKNOWLEDGEMENTS

Z.J. was supported by the National Institutes of Health under Award Number U54AG075936 and by the Whitehead Scholars Program at Duke University School of Medicine.

## AUTHOR CONTRIBUTIONS

Z.J. conceived the study. Y.W., J.Z., H.X., and C.H. performed the analysis. J.Z., Z.T., D.Z., G.T., D.L., and Z.J. provided computational resource and advised the analysis. Y.W., J.Z., H.X., D.L, and Z.J. wrote the manuscript.

## KEY POINTS

- The performance of 18 computational methods for cell segmentation was systematically evaluated.
- General-purpose methods designed with the attention mechanism show the best overall performance.
- Various factors, including image channels, choice of training data, and cell morphology may affect segmentation performance.
- Seggal is an online resource for downloading pre-trained segmentation models.

