## Supplementary Figure for "A systematic evaluation of computational methods for cell segmentation"

### METHODS

#### Datasets

**TissueNet**[1] dataset V1.1, obtained from the TissueNet website[2], comprises 7,022 fluorescence staining images. In the original dataset, the images are distributed as 2,580 images in the training set, 3,118 in the validation set, and 1,324 in the testing set. The images span ten tissue types of breast, colon, epidermis, esophagus, lung, lymph node, pancreas, spleen, tonsil, and lymph node metastasis, and originate from three species of human, mouse, and macaque. Each image features a nuclear staining channel and a membrane or cytoplasm whole-cell staining channel, along with annotations for both nuclei and whole cells, derived from an iterative human-in-the-loop process[1]. The training set images have a resolution of $512 \times 512$ pixels, while those in the validation and testing sets are $256 \times 256$ pixels each.

**LIVECell**[3] dataset was obtained from the LIVECell website[4]. The original dataset encompasses 5,387 microscopy images, divided into 3,253 for training, 570 for validation, and 1,564 for testing. These images represent eight distinct cell types with unique morphological features, including A172, BT-474, BV-2, Huh7, MCF7, SH-SY5Y, SkBr3, and SK-OV-3. Each image, with a resolution of $704 \times 520$ pixels, includes manually labeled and expert-validated cell annotations.

**Imaging Mass Cytometry (IMC)** images were obtained from the `imcdatasets` R/Bioconductor package. All five datasets included in the `imcdatasets` package were included: pancreas, melanoma, cancer, breast cancer, and spheroids. One image was randomly selected for each dataset. For each image, the "DNA1" channel was used as a single-channel image.

**10x Visium** images were downloaded from 10x website (https://www.10xgenomics.com/datasets). DAPI images from five tissues were obtained: human breast cancer, human glioblastoma, human prostate, human skin melanoma, and human lung cancer.

**10x Xenium** images were downloaded from 10x website (https://www.10xgenomics.com/datasets). DAPI images from five tissues were obtained: human breast cancer, human lymph node, human lung, mouse brain coronal, and human pancreatic cancer.

**Vizgen** images were downloaded from Vizgen website (https://info.vizgen.com). DAPI images from five tissues were obtained: human colon cancer, human liver cancer, human ovarian cancer, mouse liver, and mouse brain receptor.

#### Image preprocessing

All images in the six datasets were preprocessed following a common practice in the field[1,5]. Specifically, each image along with the corresponding annotations were cropped to sliding windows. The size of each window is $256 \times 256$ and the step size is 75% of window size (192 pixels). The sliding windows scan across the image, from left to right and top to bottom, at each step making a local crop. After reaching the end of a row, the window moves down by the step size and continues scanning until the entire image has been covered, ensuring comprehensive coverage and increasing the likelihood of accurate detection. For images from IMC, 10x Visium, 10x Xenium, and Vizgen, we selected a subset of cropped images such that the number of cells included in the selected images exceeds 100 for each tissue and each technology platform. Pixel values were then normalized to account for exceptionally bright, isolated pixels[6]. Specifically, the pixels values were first subtracted by $L$ and then divided by $U - L$. Here $L$ and $U$ are the 0.1% and 99.9% percentiles of pixel values in the original image, respectively. Furthermore, we employed Contrast Limited Adaptive Histogram Equalization (CLAHE)[7] to balance the dynamic range within each image.

For images from 10x Visium, 10x Xenium, and Vizgen datasets, we additionally performed downsampling by using the "resize" function provided by OpenCV, where interpolation parameter was set as bilinear interpolation.

#### Details of segmentation methods

Unless otherwise specified in the following, all methods were trained and applied with the default parameters. Each method was adjusted to fit the input image dimensions, including height, width, and channel number. The training and validation sets from the TissueNet and LIVECell datasets were used as inputs for each method during model training. Subsequently, the trained model was applied to the testing set for cell segmentation and performance evaluation. Each method was independently executed on a dataset for a specific segmentation task. For each task, we allocated 200G of memory and either a single GPU or multiple GPUs of the same type, depending on the method's support for multi-GPU training. MMDetection (version 2.28.2), an open-source object detection toolbox, was used to implement the following methods: Swin-S, Swin-T, Cascade Mask RCNN seesaw, SOLOv2, Res2Net, RF-Next, HRNet, Mask2former, Mask RCNN, and MS RCNN. The methods benchmarked in this study are described as follows.

**Cellpose**[5] employs the general U-Net[8] architecture, which involves multiple downsampling steps of convolutional features, followed by a symmetrical upsampling process. Instead of feature concatenation, Cellpose uses direct summation between the upsampling and downsampling phases to reduce the number of parameters. Additionally, its standard U-Net building blocks are replaced with residual blocks[9] to enhance performance. Cellpose (version 1.0.2) was downloaded from https://github.com/MouseLand/cellpose and trained on a single GPU.

**Mesmer**[1] utilizes a ResNet-50[9] backbone model for feature pyramid extraction in image analysis. These extracted features are subsequently transformed into pyramid structures via a Feature Pyramid Network (FPN)[10]. The pyramid features are then employed to generate predicted cell instance masks, matching the size of the input image, through semantic segmentation heads. Mesmer (version 0.12.9) was downloaded from https://github.com/vanvalenlab/deepcell-tf. For training, Mesmer was configured to run on 4 GPUs with a learning rate of 0.0004, a batch size of 8, and a total of 200 epochs.

**StarDist**[11] employs star-convex polygons for cell instance localization, offering superior shape representation compared to bounding boxes without the need for further shape refinement. It is built upon the U-Net[8] architecture. To prevent feature competition between the two subsequent output layers, StarDist integrates an additional $3 \times 3$ convolutional layer with 128 channels, accompanied by ReLU activations. It utilizes a single-channel convolutional layer with sigmoid activation for object probability output, and for polygon distance estimation, the number of output layers aligns with the radial directions without an extra activation function. StarDist (version 0.8.3) was downloaded from https://github.com/stardist/stardist. StarDist was trained on a single GPU for 100 epochs.

**RetinaMask**[12], an advanced version of RetinaNet[13], serves as a unified, one-stage object detection system. It utilizes a ResNet-50 backbone network to compute a comprehensive convolutional feature map. This backbone connects to two subnetworks: one for convolutional object classification and another for bounding box regression, based on the extracted feature map. The key advancement of RetinaMask over RetinaNet is the introduction of instance mask prediction during training. As a result, RetinaMask outputs not only bounding boxes and classifications but also precise cell instance masks. RetinaMask (version 0.1.1) was downloaded from https://github.com/vanvalenlab/deepcell-retinamask. For training, RetinaMask was configured to run on 4 GPUs with a learning rate of 0.0005 over 100 epochs.

**FeatureNet**[14] consists of an encoder for generating a low-dimensional feature representation of the input image, utilizing convolutional filters, activation functions, and max-pooling operations. These features are then input into a classifier, based on a fully connected network, which assigns probability scores to three classes: boundary, interior, and background. These probabilities are used to create the final cell instance masks. FeatureNet (version 0.12.9) was downloaded from https://github.com/vanvalenlab/deepcell-tf. For training, FeatureNet was configured to run on 4 GPUs with a learning rate of 0.001, a batch size of 16, and 25 epochs.

**Centermask2**[15] comprises three key components. The first is the VoVNetV2, an efficient backbone network for feature extraction. The second, FCOS[16], is an anchor-free, proposal-free object detection network that directly predicts bounding box coordinates through per-pixel analysis. The final component, the Spatial Attention-Guided Mask (SAG-Mask), utilizes outputs from the first two components for precise cell instance mask prediction. Centermask2 was downloaded from https://github.com/youngwanLEE/centermask2. Centermask2 was configured to run on 4 GPUs with 100,000 iterations and a batch size of 32.

**Mask RCNN**[17] follows the spirit of Faster R-CNN architecture[18] with the same two-stage procedure. The first stage generates proposals about the regions where there is an object (Region Proposal Network, RPN[18]), which is identical first stage with Faster R-CNN. In the second stage, in parallel to the branch predicting the class and bounding box, Mask RCNN adds an additional segmentation branch to output a binary mask for each Region of Interest (ROI). Mask RCNN was configured to run on 4 GPUs with 100 epochs and 24 batch sizes.

**MS RCNN** (Mask Scoring R-CNN)[19] improves upon the instance segmentation capabilities of Mask RCNN[20]. Unlike Mask RCNN, which uses the segmentation branch's output as the final mask prediction, MS RCNN adds a mask scoring branch. This branch calculates a mask $IoU$ (Intersection over Union) score, assessing the overlap between the predicted and ground truth binary masks. This score provides a more accurate evaluation of instance segmentation quality without adding significant complexity or computational burden. MS RCNN was trained on 4 GPUs with a configuration of 50 epochs and a batch size of 48.

**ResNeSt**[21] introduces a multi-branch architecture that employs channel-wise attention across its branches, combining the strengths of feature-map attention and multi-path representation. Its key innovation, the Split-Attention (SA) module, acts as a modular, versatile block, easily substituting the conventional residual block. This module fosters more diverse representations through cross-feature interactions. Such diversity enhances segmentation performance without increasing model complexity or reducing computational efficiency, thereby offering a scalable solution for various tasks. ResNeSt (version 0.1.1) was downloaded from `https://github.com/chongruo/detectron2-ResNeSt`. ResNeSt was configured to run on 4 GPUs with a ResNet FPN backbone, performing 50,000 iterations and using a batch size of 4.

**Mask2former**[22], a universal image segmentation architecture, evolves from the Maskformer meta-architecture[23] and includes a ResNet-50 backbone for feature extraction, a pixel decoder, and a Transformer decoder. Its key advancement is the adoption of a Transformer decoder that applies masked attention, focusing on localized features around predicted segments, rather than using conventional cross-attention. These segments, derived from the pixel decoder, enable the refined output of object instances. Mask2former was configured to run on 4 GPUs for 50 epochs and a batch size of 36.

**Cascade Mask RCNN seesaw**[24] enhances Cascade Mask RCNN by introducing an innovative seesaw loss function to address the class imbalance issue common in object detection and instance segmentation tasks. While traditional methods like Focal Loss[13] provide some relief, they may fall short in extreme class imbalance situations. Seesaw loss dynamically adjusts loss weights for each class, based on the class sample size in each mini-batch. It employs a seesaw mechanism that increases the weight for minority classes and decreases it for majority classes, ensuring more balanced learning. Additionally, it gives more weight to harder samples within each class to further equalize the learning process. Cascade Mask RCNN seesaw was configured to run on 4 GPUs for 50 epochs and a batch size of 8.

**Swin Transformer**[25] merges the strengths of convolutional neural networks (CNNs) and transformers, offering a scalable and efficient approach for various visual tasks. Its hierarchical structure is a key innovation, utilizing shifted window-based self-attention mechanisms to process local information within small, discrete windows. These windows are progressively merged and resized across stages, enabling the model to effectively capture both local and global contexts. The primary distinction between Swin-T and Swin-S lies in their size and computational complexity, with Swin-S being twice as large as Swin-T. Both variants were trained on 4 GPUs for 50 epochs and a batch size of 8.

**SOLOv2**[26], an evolution of the original SOLO[27], adopts a refined approach that focuses on direct per-pixel predictions instead of conventional bounding-box techniques. At its core, SOLOv2 features a fully convolutional network structure, category-specific kernel predictions, and the innovative Matrix Non-Maximum Suppression (Matrix NMS) for refining overlapping instances[28]. Initially, it utilizes the same ResNet-50 backbone and FPN as SOLO for feature extraction, then integrates a category-specific kernel prediction network to infer all instance masks. Matrix NMS, notably faster and more accurate than traditional NMS[29], enhances the accuracy and efficiency of the model. SOLOv2 was trained on 4 GPUs for 60 epochs with a batch size of 8.

**RF-Next**[30] primarily follows the Multi-Stage Temporal Convolutional Network (MS-TCN)[31] approach, introducing a global-to-local search method that includes two innovative components: a genetic-based global search algorithm for generating competitive receptive field combinations, and an expectation-guided iterative local search for fine-tuning these combinations. Specifically, RF-Next uses Mask RCNN as its instance segmentation foundation and integrates these components into convolutional layers with kernels larger than one, optimizing segmentation results. RF-Next was configured to run on 4 GPUs for 200 epochs with a batch size of 8.

**HRNet**[32] introduces an architecture that features parallel multi-resolution convolutions (PMRC), repeated multi-resolution fusions (RMRF), and a representation head maintaining high-resolution representations throughout. Diverging from traditional models that downsample and then upsample, HRNet starts with a high-resolution subnetwork and progressively integrates additional high-to-low resolution subnetworks in stages. It continuously fuses these multi-resolution subnetworks in parallel, enabling dynamic feature interplay. HRNet's core design focuses on iterative multi-resolution fusions, constantly exchanging and integrating information across these parallel subnetworks to capture both fine and coarse features in the input data comprehensively. HRNet was configured to run on 4 GPUs for 200 epochs with a batch size of 24.

**Res2Net**[33] introduces a modification to the ResNet architecture with a granular multi-scale design. Unlike traditional ResNet models, Res2Net splits input features in each residual block into subsets. Each subset processes different scales using varying dilation rates, functioning similarly to parallel sub-ResNets. These multi-scale feature maps are then efficiently integrated through convolution, enabling effective information sharing between scales. The key strength of Res2Net lies in its hierarchical block structure, which facilitates enhanced feature propagation and efficient exploitation of multi-scale features. Res2Net was configured to run on 4 GPUs for 20 epochs with a batch size of 8.

**Segment Anything**[34] introduces the Segment Anything Model (SAM) for promptable segmentation, incorporating three key components: a Masked AutoEncoders (MAE)[35,36]-based image encoder adapted for high-resolution inputs, a flexible prompt encoder (PE) capable of processing various prompts, and a fast mask decoder (ME). SAM utilizes a pre-trained Vision Transformer (ViT[37]) as its image encoder. The prompt encoder can handle both sparse prompts (interpreted via positional encodings[38–40] and combined with specialized embeddings) and dense prompts (integrated with the encoded image). The mask decoder then amalgamates the embeddings from both components to generate segmentation masks using a modified Transformer decoder block[41], followed by a dynamic mask prediction head, which outputs the mask foreground probability for each pixel. Segment Anything, downloaded from `https://github.com/facebookresearch/segment-anything`, was employed for segmentation, utilizing its default settings and the pre-trained ViT-H SAM model. Consequently, the process did not involve any GPU-based training.

#### Segmentation benchmarks

We assessed segmentation performance in this study using the `pycocotools` Python package (version 2.0.6) that implements the COCO evaluation metrics[1,42,43] , a standard evaluation method in instance segmentation. To ensure all cell instances per image were considered, we increased the parameter for maximum detections per image to 3000, from the default value of 100. The details of the evaluation metrics are described below:

##### *Intersection over Union ($IoU$)*

For the $o$th image in the test set with resolution of $H \times W$ pixels, let $\{S^p_{k,o} \in \{0,1\}^{H\times W}\}^K_{k=1}$ be the predicted instances of $K$ cells, and let $\{S^g_{k',o} \in \{0,1\}^{H\times W}\}^{K'}_{k'=1}$ be the ground truth annotations of $K'$ cells. The Intersection over Union ($IoU$) score of the $k$th predicted cell instance is computed by:

$$IoU^p_{k,o} = \max\left(\left\{\frac{S^p_{k,o} \cap S^g_{k',o}}{S^p_{k,o} \cup S^g_{k',o}}\right\}^{K'}_{k'=1}\right). \tag{1}$$

Likewise, the $IoU$ score of the $k'$th ground truth cell annotation is computed by:

$$IoU^g_{k',o} = \max\left(\left\{\frac{S^p_{k,o} \cap S^g_{k',o}}{S^p_{k,o} \cup S^g_{k',o}}\right\}^{K}_{k=1}\right). \tag{2}$$

##### *Precision and Recall*

Let $t$ be a threshold taking values from $\{t \mid t = 0.5+0.05i, 0 \le i \le 9\}$. For a given threshold $t$, a predicted cell instance $k$ is considered a true positive (TP) if $IoU^p_{k,o} \ge t$, and a false positive (FP) if $IoU^p_{k,o} < t$. A ground truth cell annotation $k'$ is considered a false negative (FN) if $IoU^g_{k',o} < t$.

For the $o$th image and with threshold $t$, let $TP_{t,o}$, $FP_{t,o}$, and $FN_{t,o}$ be the total number of true positive, false positives and false negatives, respectively. The precision $P_{t,o}$ is defined as $P_{t,o} = \frac{TP_{t,o}}{TP_{t,o}+FP_{t,o}}$. The recall $R_{t,o}$ is defined as $R_{t,o} = \frac{TP_{t,o}}{TP_{t,o}+FN_{t,o}}$.

##### *Average Precision and Average Recall*

Average precision $AP_t$ was calculated as the area under the precision-recall curve. This curve, which is monotonically decreasing, was constructed using the calculated precision $\{P_{t,o}\}^O_{o=1}$ and recall $\{R_{t,o}\}^O_{o=1}$ across all images. Average Recall $AR_t$ was calculated as the average of recall $\{R_{t,o}\}^O_{o=1}$ across all images. Mean average precision $mAP$ was calculated as the average of $AP_t$ across all possible values of $t$.

#### Cell morphology

**Cell convexity.** Let $r_{\text{cell}}$ be the perimeter of a cell's boundary, and let $r_{\text{convex}}$ be the perimeter of the cell's convex hull, the smallest possible convex shape that completely contains the cell region. The cell convexity of the cell is defined as $r_{\text{convex}}/r_{\text{cell}}$.
**Cell elongation.** Let $B$ be a set of two-dimensional points representing the boundary of a cell. We compute $e_a$ and $e_b$, the two eigenvalues of the covariance matrix of $B$, where $e_a > e_b$. Cell elongation is computed as $e_b/e_a$.
**Cell size.** The size of a cell instance is defined as the number of pixels contained within the boundary of the cell.

#### Multiple linear regression for mAP

For each tissue or cell type, we calculated the average cell size and cell elongation across all cells and images.

In TissueNet, we performed min-max scaling on the number of training images, average cell size, and average cell elongation across all tissues to ensure comparability in scale. Let $m$, $N$, $S$, and $E$ represent the mean average precision (mAP), scaled number of training images, scaled average cell size, and scaled average cell elongation, respectively. We then fitted the following regression model.

$m \sim N + S + E$

In LIVECell, since the number of training images is identical for all cell types, it was not included in the regression analysis. Similar to the approach in TissueNet, min-max scaling was applied to the average cell size and average cell elongation across all cell types. We fitted the following regression model using the same notation as in TissueNet.

$m \sim S + E$

#### Morphological Operations

The `binary_dilation` and `binary_erosion` functions from the `scipy` (version 1.9.2) multidimensional image processing (`ndimage`) package were used for cell nuclei mask dilation and whole cell mask erosion, respectively. The dilation process stopped for a cell when its boundary reached the boundaries of other neighboring cells.

#### Scalability

For each dataset evaluated in this study, we recorded the running time $T$ of a segmentation method as the time taken to finish model training after all packages were loaded. Different segmentation methods have different choices of batch size and number of epochs or iterations, which may affect their running time. To ensure a fair comparison across methods, we calculated standardized running time as $T_s = T \times \frac{B}{8} \times \frac{50}{E}$ for methods using epochs as a parameter, and $T_s = T \times \frac{B}{8} \times \frac{50N}{I \times B}$ for methods using iteration as a parameter. Here $B$, $E$, $I$, and $N$ represent the batch size, number of epochs, number of iterations, and number of training images, respectively.

#### Usability

We evaluated the usability of each method using three metrics, each manually scored on a scale from 1 to 10, with higher scores indicating better performance. The scores for these three metrics are shown in Supplementary Table 1.

**Code Maintenance** assesses the ease of updating and maintaining a method's programming code. This metric takes into account the clarity and readability of the code, adherence to good software engineering practices (such as modular design, encapsulation, version control system usage, and comprehensive commenting), and the availability of supporting documentation.

**Ease of Use** assesses the simplicity with which end users and developers can utilize and interact with the method. Key considerations include the ease of training the model, adjusting parameters, interpreting outputs, and resolving any encountered issues.

**Hardware Support** assesses a method's compatibility with various GPU architectures and its ability to fully leverage specific hardware features, such as GPUs' parallel processing capabilities. Additionally, this criterion considers the method's adaptability to different hardware configurations and its capacity to efficiently scale according to available resources.

#### Ranking Scheme

The methods are ranked by their overall performance. The overall performance is calculated as the average of seven metrics: the accuracy in TissueNet nuclei segmentation with dual-channel images, the accuracy in TissueNet nuclei segmentation with nuclei images, the accuracy in TissueNet whole cell segmentation with dual-channel images, the accuracy in TissueNet whole cell segmentation with whole cell images, the accuracy in LIVECell whole cell segmentation, averaged usability score, and averaged scalability score.

For each of the five segmentation tasks, the accuracy scores is calculated as the average of five metrics: $mAP$, $AP_{0.5}$, $AP_{0.75}$, $AR_{0.5}$, and $AR_{0.75}$.

The averaged usability score is calculated as the average of normalized scores of code maintenance, ease of use, and hardware support. To obtain the normalized score of code maintenance, the raw scores of code maintenance across 18 segmentation methods were first scaled to have a mean of 0 and a standard deviation of 1. The normalized score was then obtained by mapping the scaled scores to a range between 0 and 1 using the cumulative density function of a standard normal distribution. Normalized scores of ease of use and hardware support were obtained similarly.

The averaged scalability score is calculated as one minus the average of normalized running time of TissueNet nuclei segmentation, TissueNet whole cell segmentation, and LIVECell segmentation. Running time of TissueNet nuclei segmentation is calculated as the average running time of TissueNet nuclei segmentation with nuclei and dual-channel images. Similarly, running time of TissueNet whole cell segmentation is calculated as the average running time of TissueNet whole cell segmentation with whole cell and dual-channel images. The three raw running time were converted to scaled and then normalized running time using the same normalization method for calculating normalized usability scores.

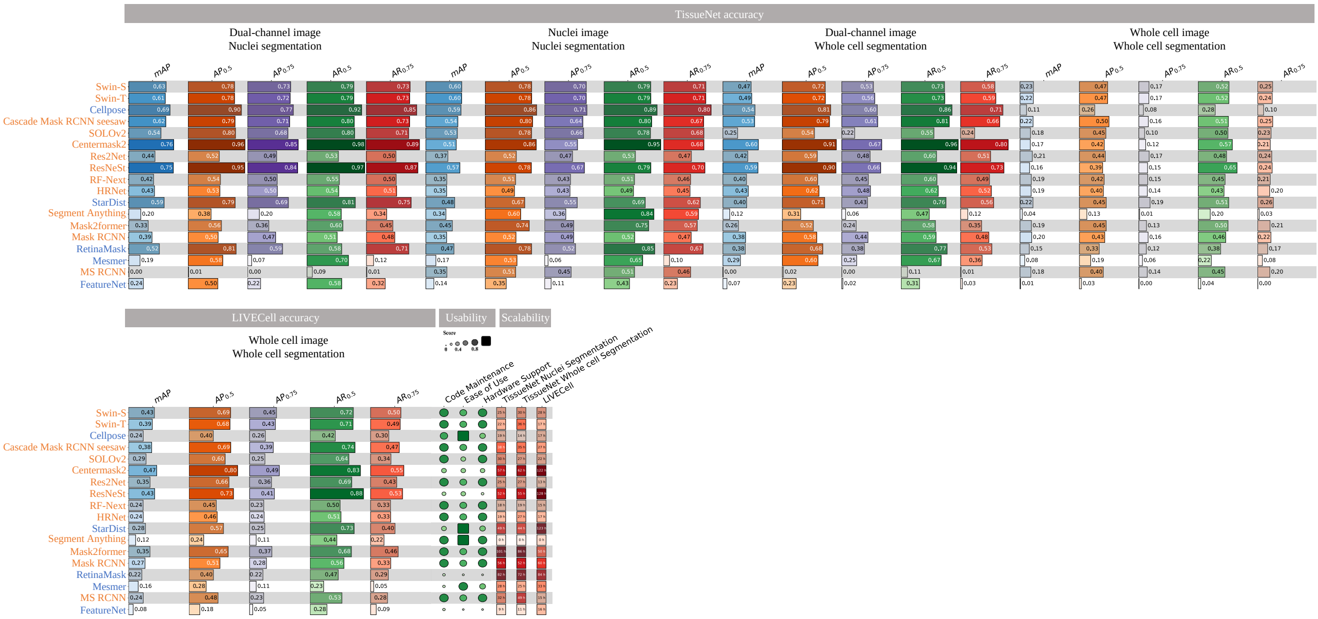


**Supplementary Figure 1.** Detailed subcategory metric scores used to calculate aggregated accuracy, usability, and scalability scores.

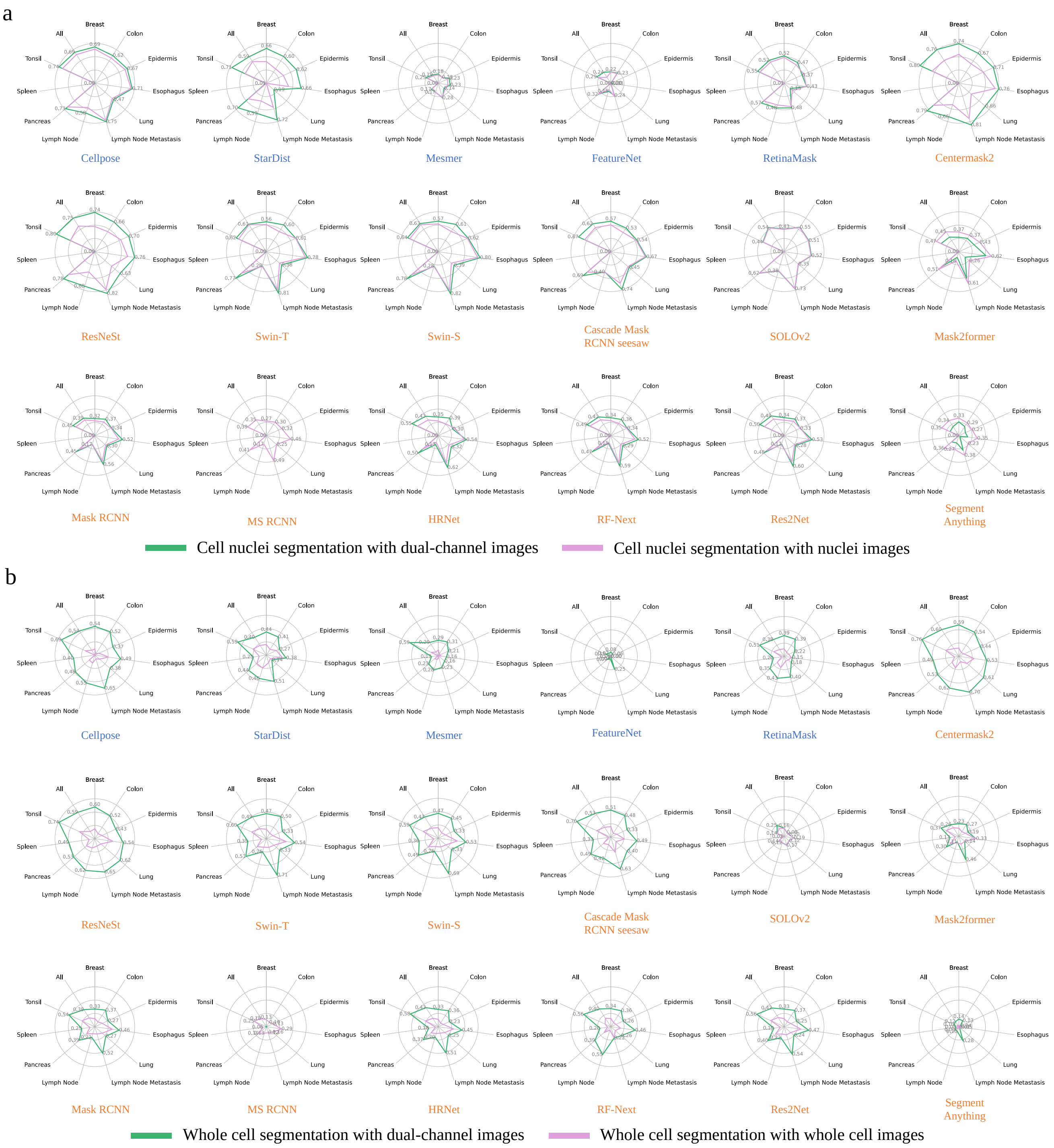


**Supplementary Figure 2.** Comparison of segmentation performances with single-channel vs dual-channel images in cell nuclei segmentation (**a**) and whole cell segmentation (**b**).

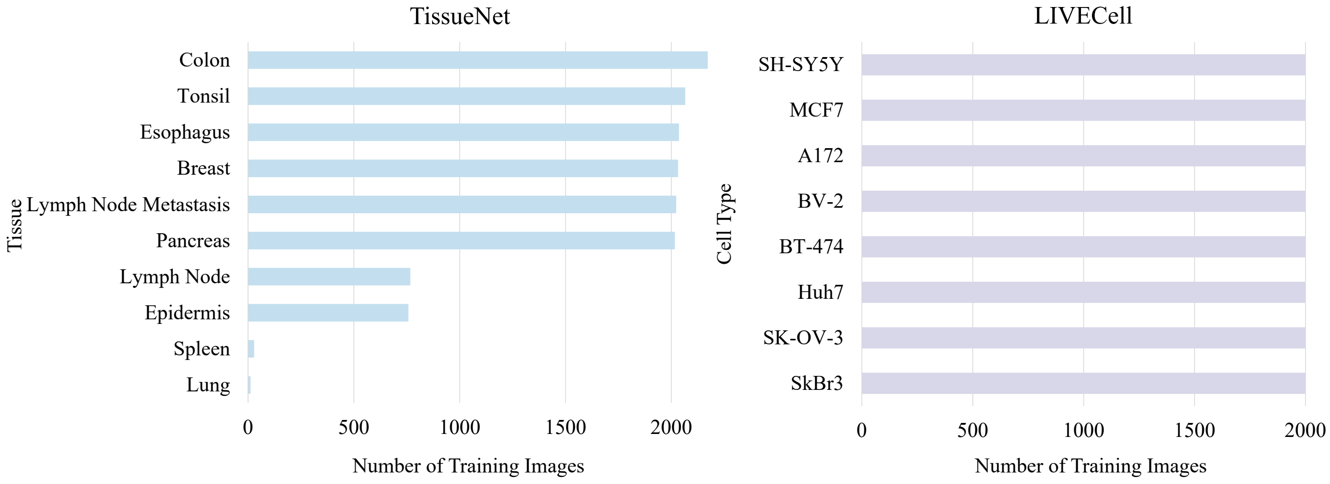


**Supplementary Figure 3.** Number of training images in TissueNet and LIVECell.

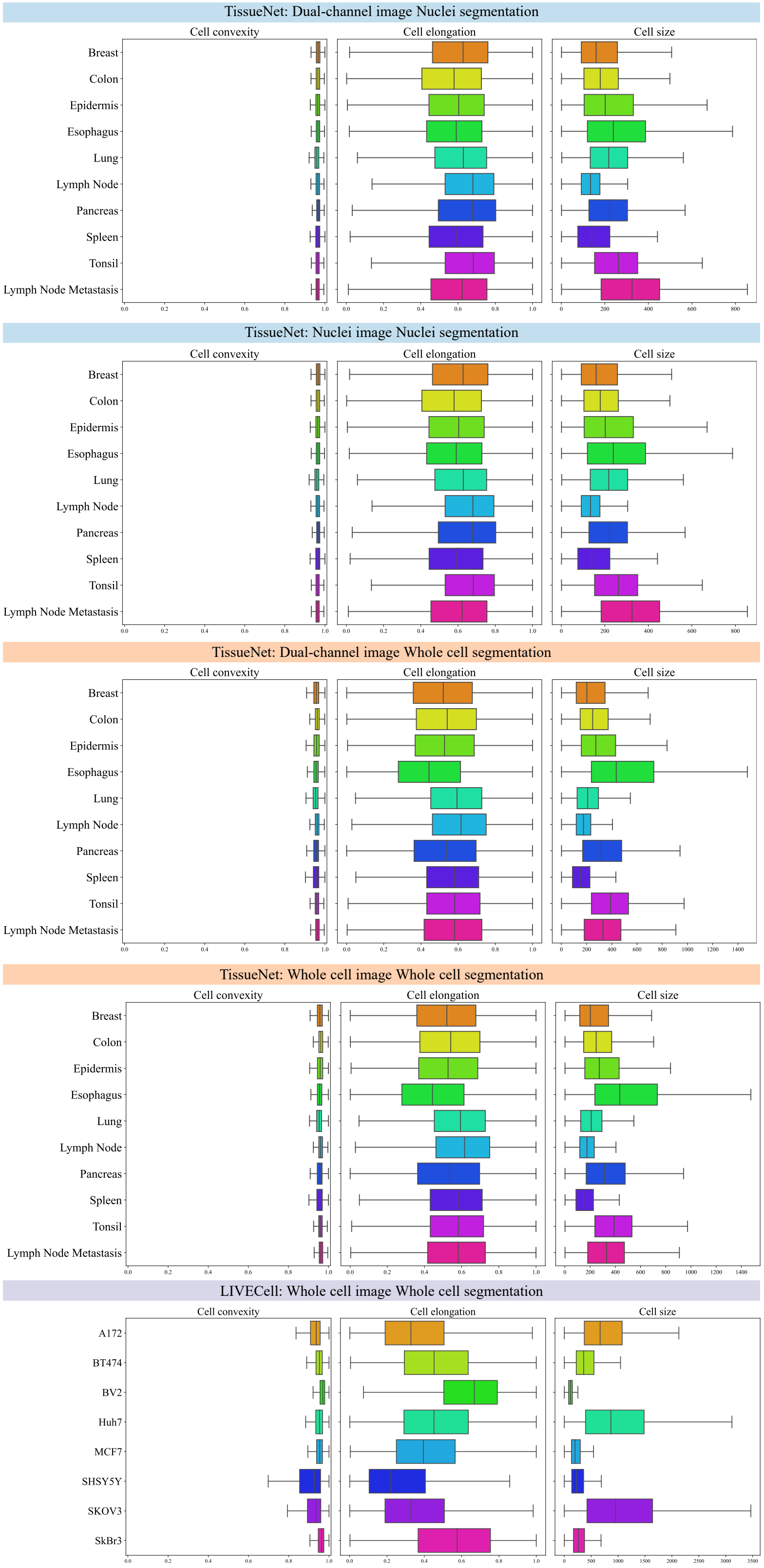


**Supplementary Figure 4.** Distributions of cell convexity, cell elongation, and cell size for each TissueNet tissue and LIVECell cell type.

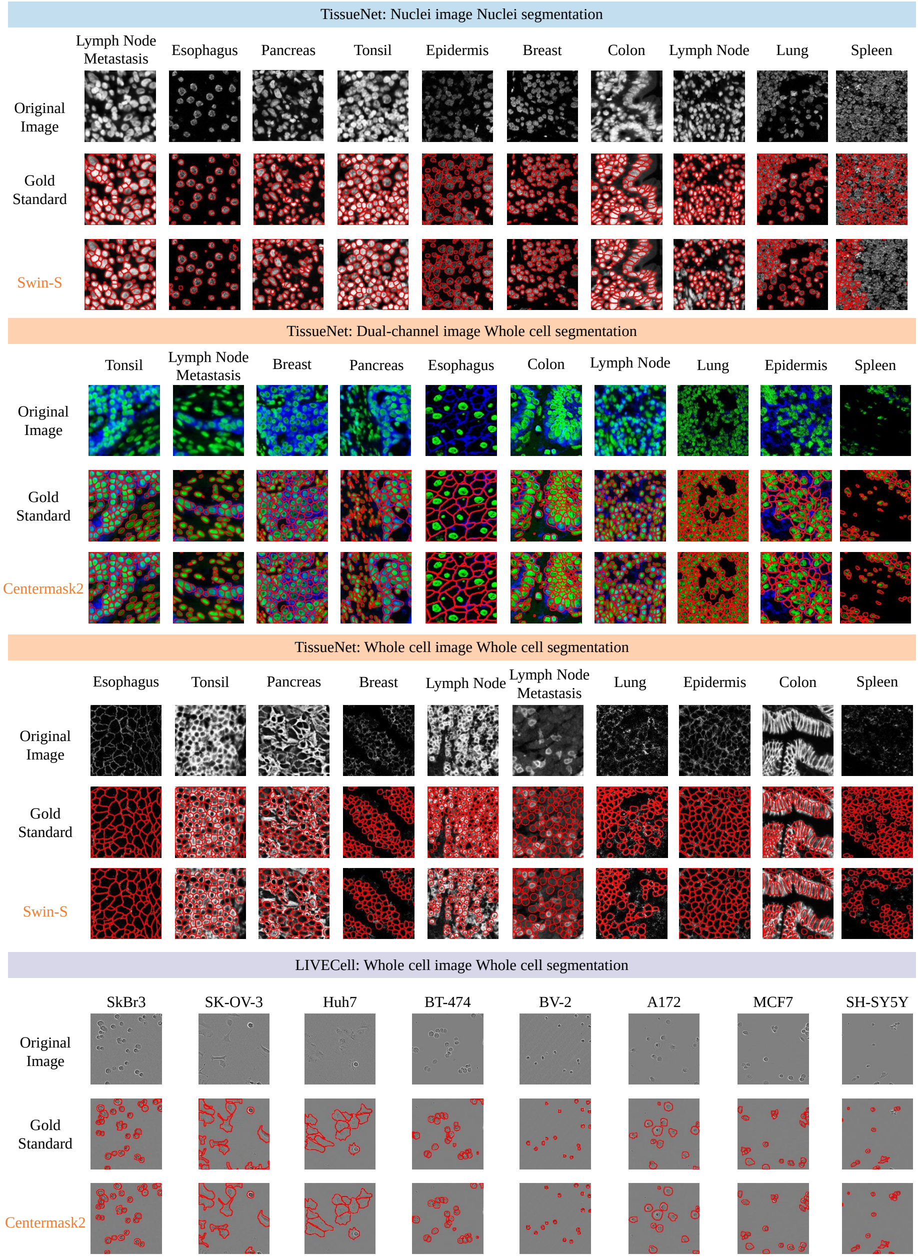


**Supplementary Figure 5.** Examples of original images, gold standard cell nuclei or whole cell masks, and segmentation results for different segmentation tasks. For each segmentation task, the method with the highest accuracy is shown.

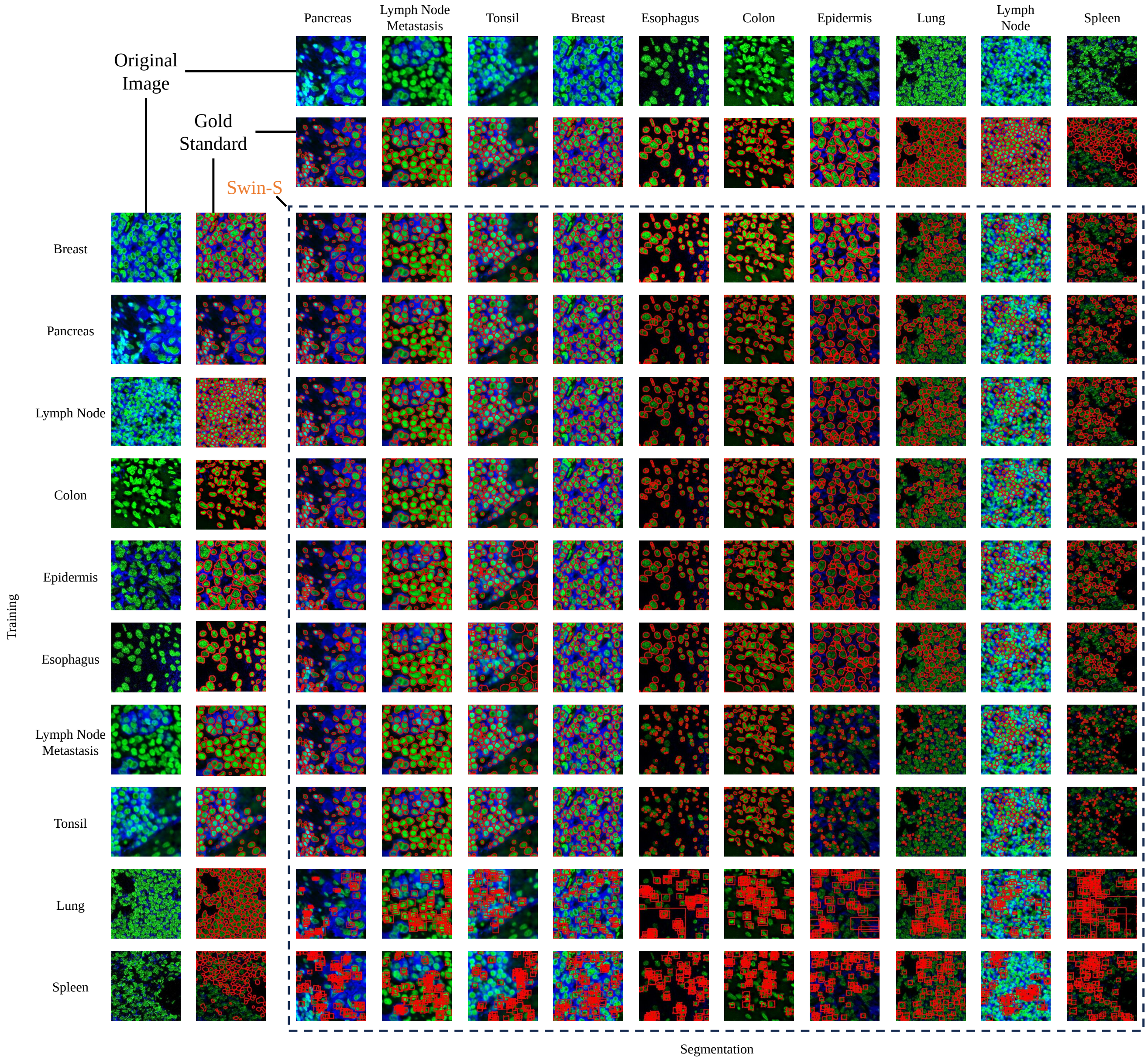


**Supplementary Figure 6.** Examples of dual-channel images from various TissueNet tissue types for training (rows) and cell nuclei segmentation (columns), including the original images, gold standard nuclei masks, and Swin-S cell nuclei segmentation results.

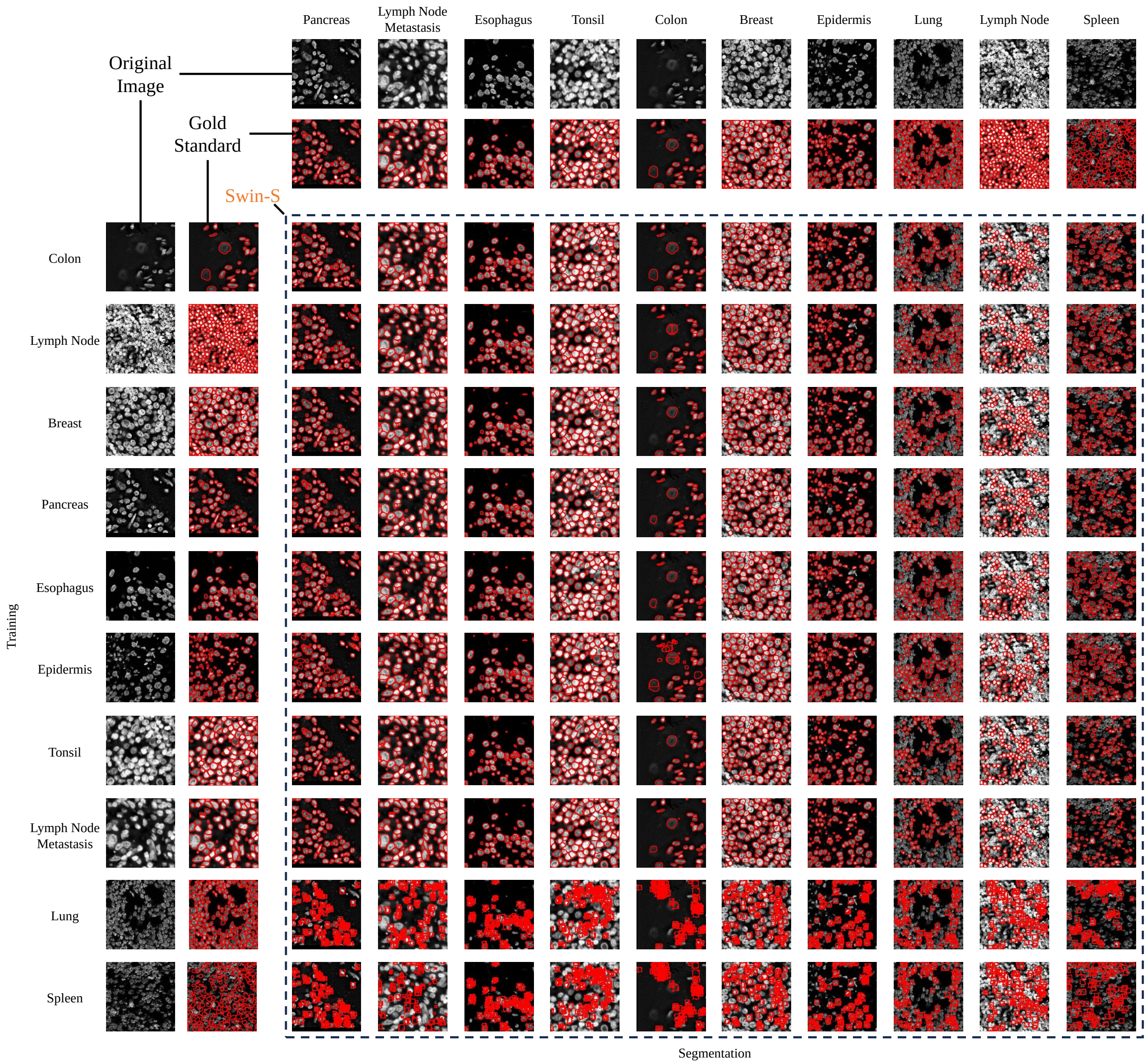


**Supplementary Figure 7.** Examples of cell nuclei images from various TissueNet tissue types for training (rows) and cell nuclei segmentation (columns), including the original images, gold standard nuclei masks, and Swin-S cell nuclei segmentation results.

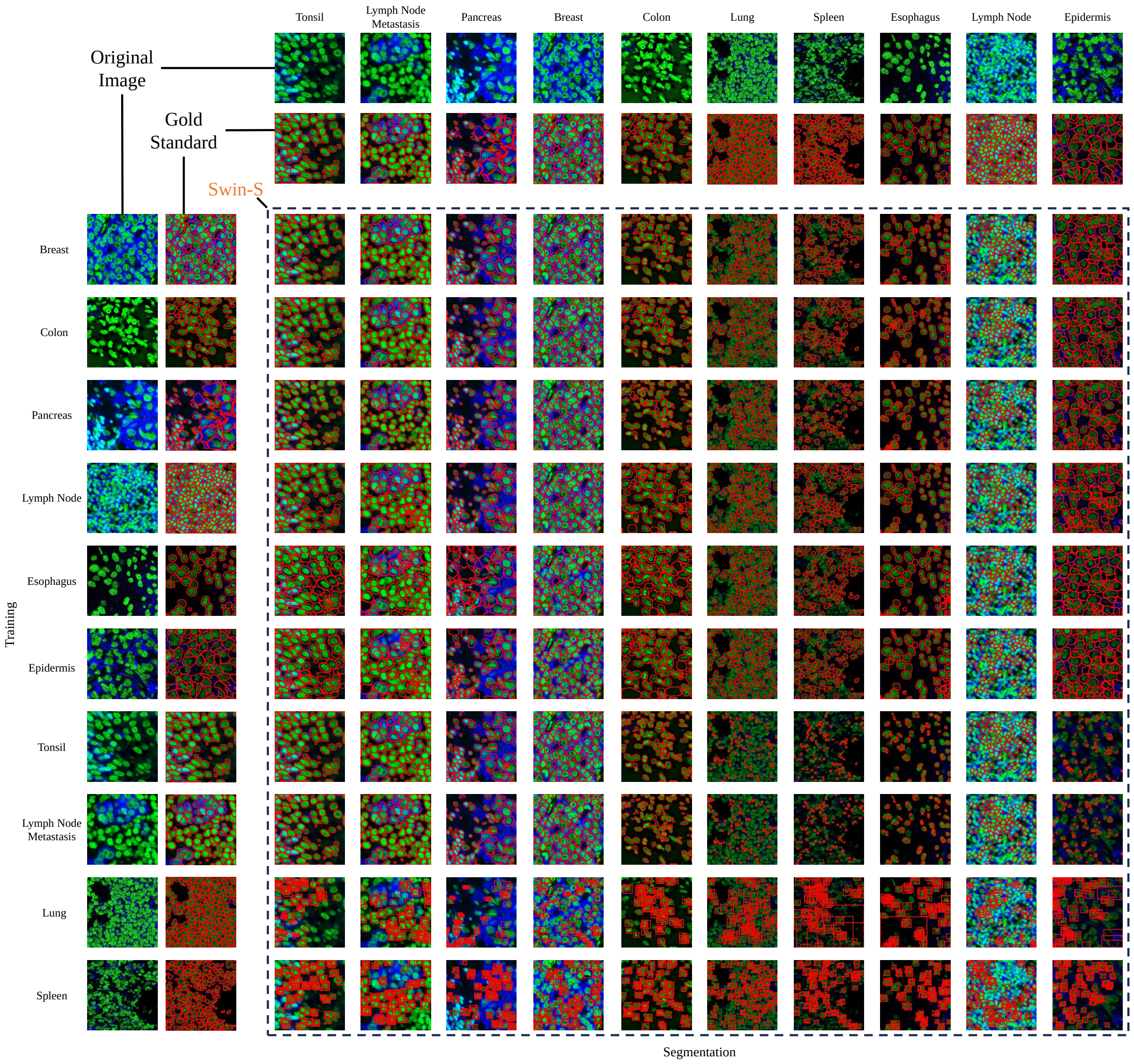


**Supplementary Figure 8.** Examples of dual-channel images from various TissueNet tissue types for training (rows) and whole cell segmentation (columns), including the original images, gold standard whole cell masks, and Swin-S whole cell segmentation results.

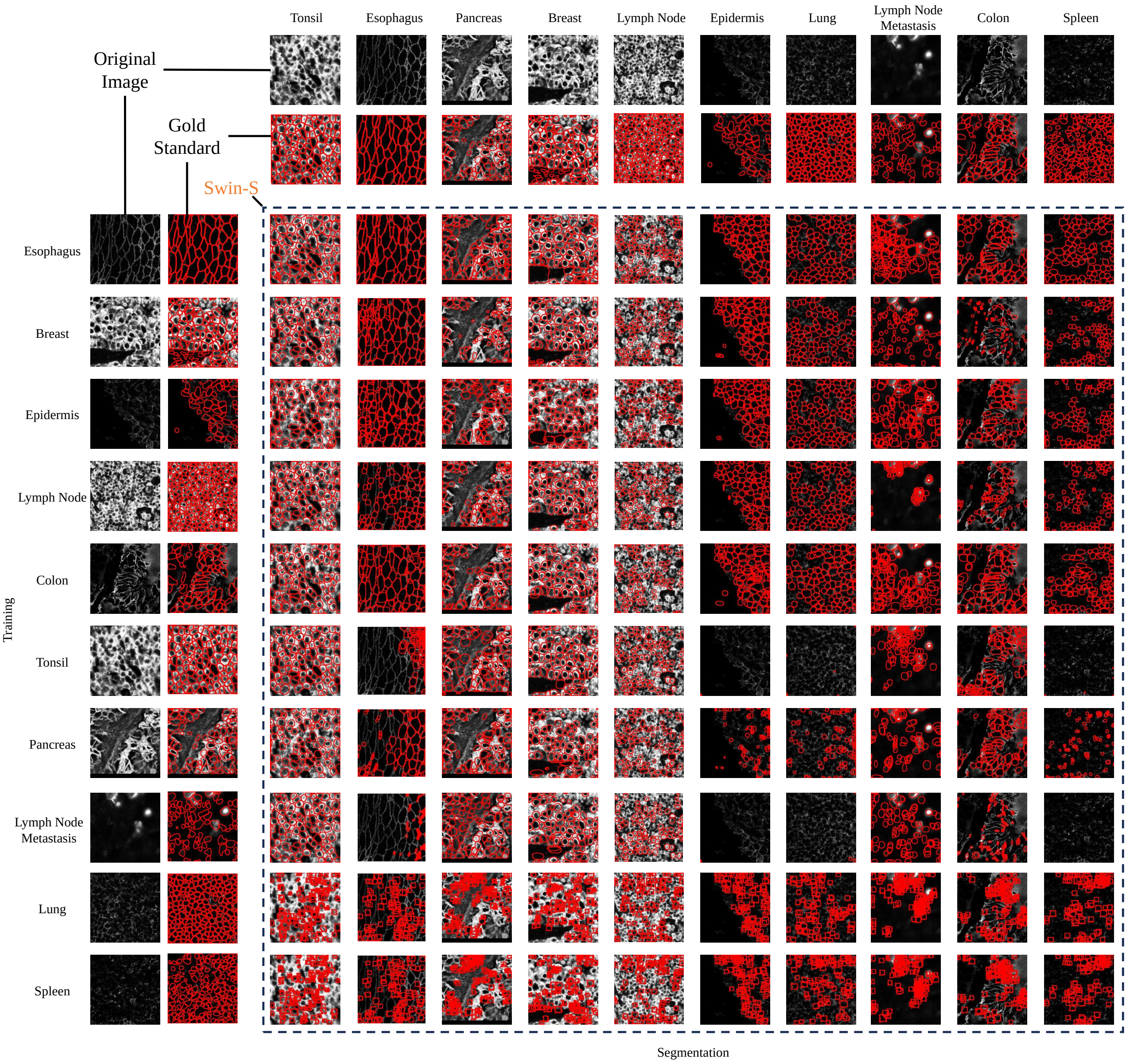


**Supplementary Figure 9.** Examples of whole cell images from various TissueNet tissue types for training (rows) and whole cell segmentation (columns), including the original images, gold standard whole cell masks, and Swin-S whole cell segmentation results.

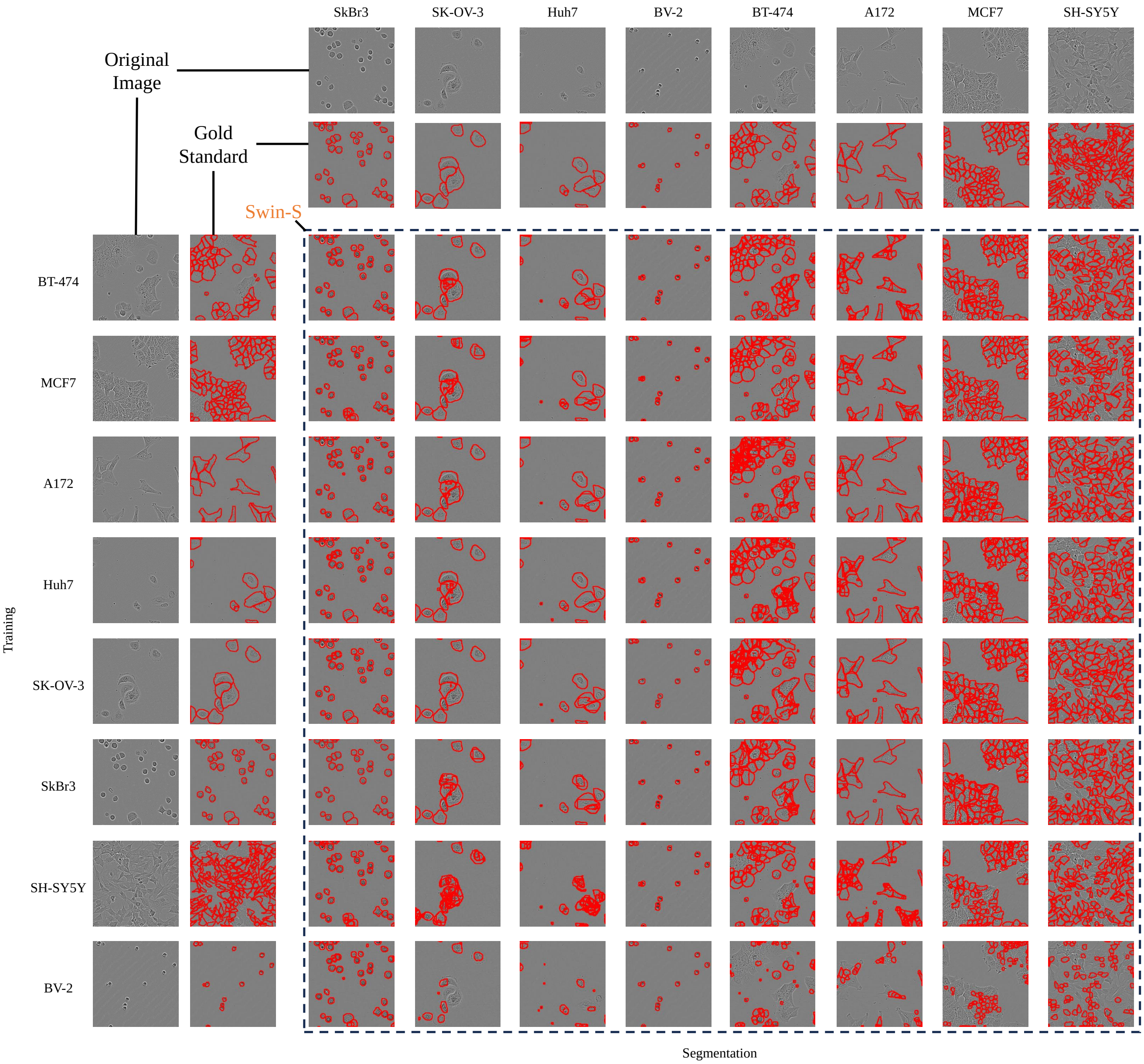


**Supplementary Figure 10.** Examples of whole cell images from various LIVECell cell types for training (rows) and whole cell segmentation (columns), including the original images, gold standard whole cell masks, and Swin-S whole cell segmentation results.

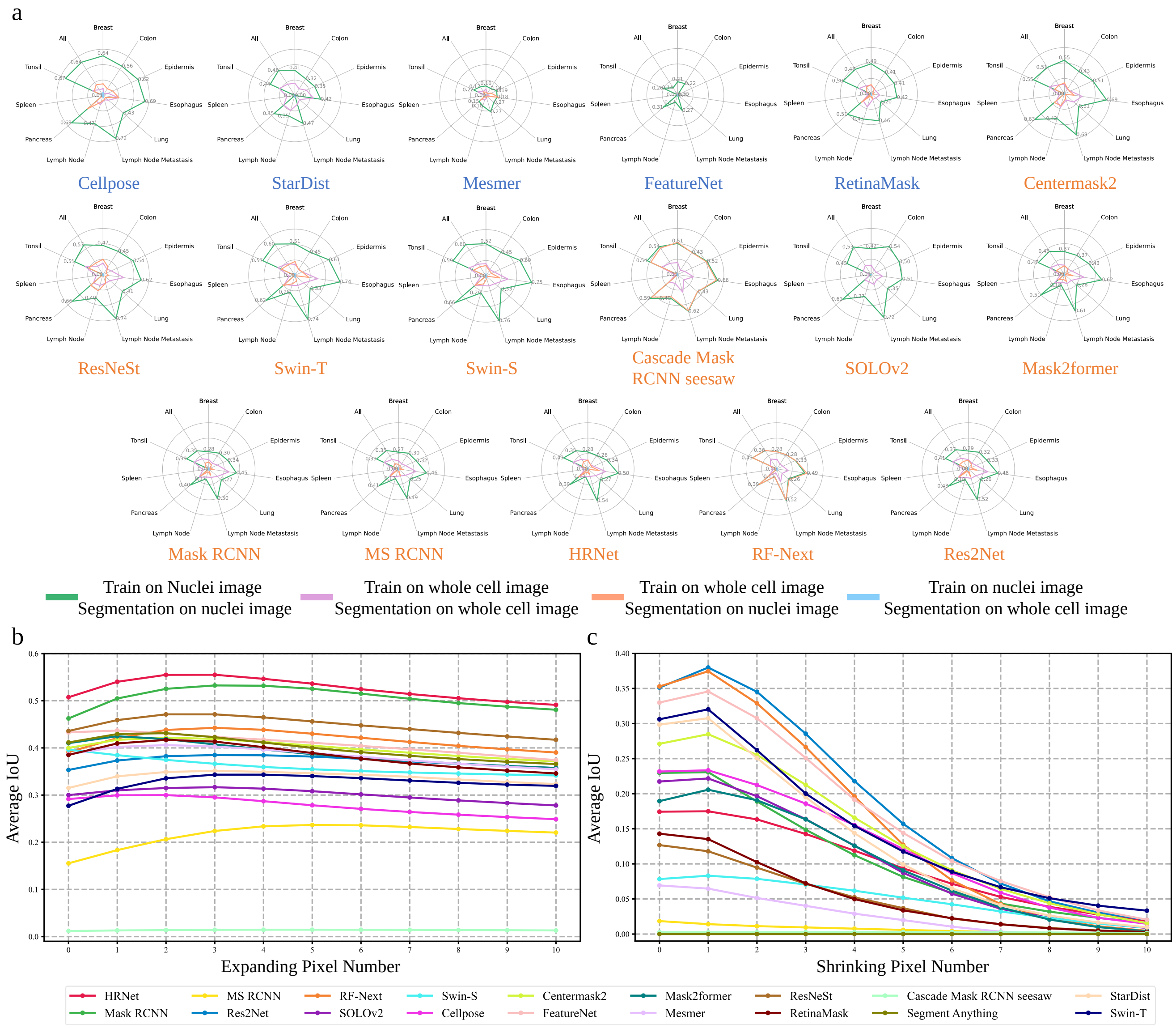


**Supplementary Figure 11.** Generalizability across image modalities. **a**, mAP for each TissueNet tissue type when training and segmentation were performed on the same or different image modalities. **b**, IoU (y-axis) between gold standard whole cell masks and expanded cell nuclei masks, averaged across all TissueNet tissue types, when cell nuclei masks were expanded by different pixel numbers (x-axis). Zero expansion represents original cell nuclei masks without expansion. **c**, IoU (y-axis) between gold standard cell nuclei masks and shrunk whole cell masks, averaged across all TissueNet tissue types, when whole cell masks were shrunk by different pixel numbers (x-axis). Zero shrinkage represents original whole cell masks without shrinkage.

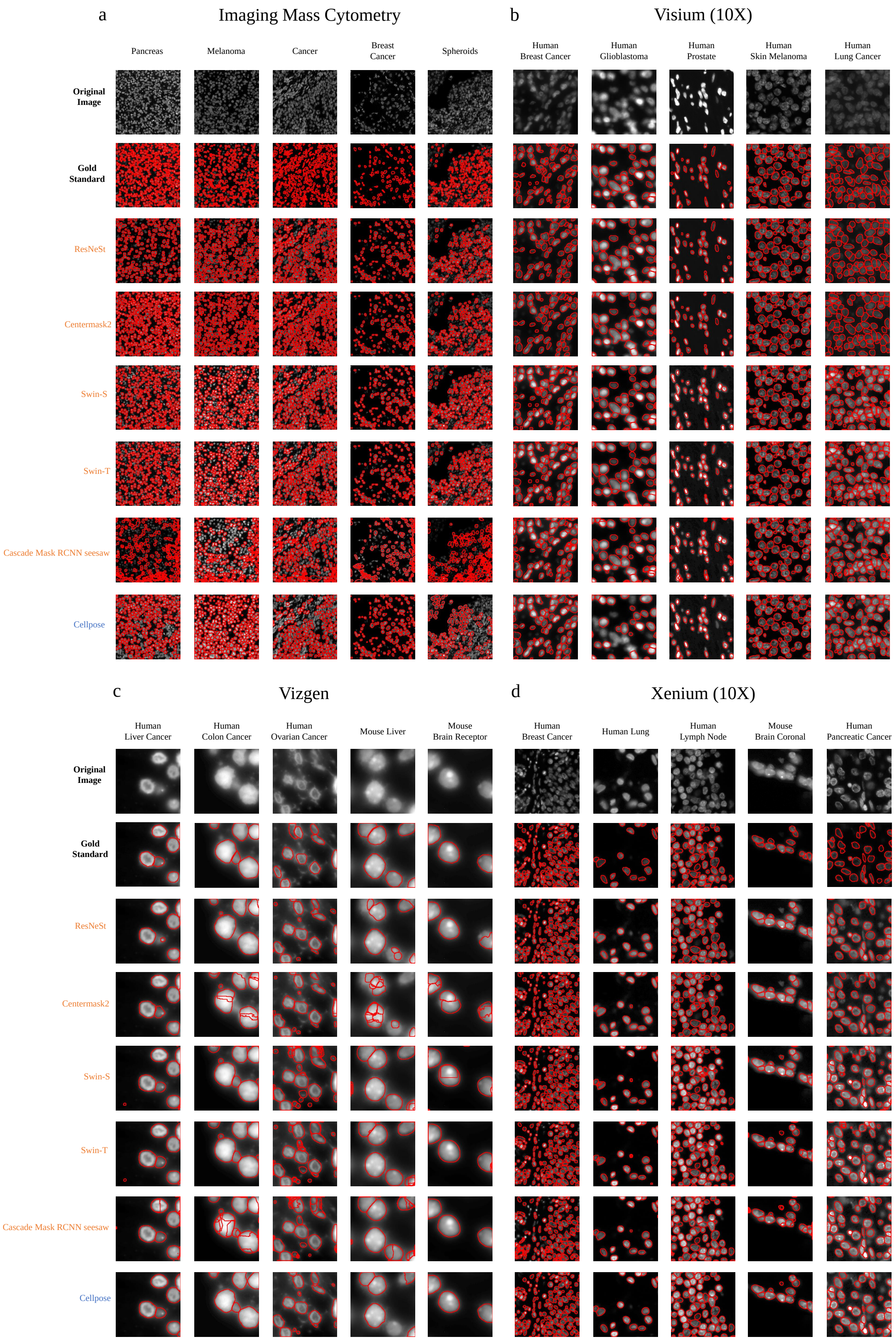


**Supplementary Figure 12.** Examples of cell nuclei images and segmentation results for different technology platforms. For each tissue and technology platform, the original image, the gold standard manually annotated nuclei boundaries, and cell nuclei segmentation from the six top-performing methods are shown. **a**, Imaging Mass Cytometry (IMC). **b**, 10x Visium. **c**, Vizgen. **d**, (10X) Xenium.
